# Response of the plant core microbiome to *Fusarium oxysporum* infection and identification of the pathobiome

**DOI:** 10.1101/2022.05.11.491565

**Authors:** Zhiguang Qiu, Jay Prakash Verma, Hongwei Liu, Juntao Wang, Bruna D Batista, Simranjit Kaur, Arthur Prudêncio de Araujo Pereira, Catriona A. Macdonald, Pankaj Trivedi, Tim Weaver, Warren C. Conaty, David T. Tissue, Brajesh K. Singh

**Author notes:** Corresponding author: Brajesh Kumar Singh.

## Abstract

Plant core microbiomes consist of persistent key members that provide critical host functions, but their assemblages can be interrupted by biotic and abiotic stresses. The pathobiome is comprised of dynamic microbial interactions in response to disease status of the host. Hence, identifying variation in the core microbiome and pathobiome can significantly advance our understanding of microbial-microbial interactions and consequences for disease progression and host functions. In this study, we combined glasshouse and field studies to analyse the soil and plant rhizosphere microbiome of cotton plants (*Gossypium hirsutum*) in the presence of a cotton-specific fungal pathogen, *Fusarium oxysporum* f. sp. *vasinfectum* (FOV). We found that FOV directly and consistently altered the rhizosphere microbiome, but the biocontrol agents enabled microbial assemblages to resist pathogenic stress. Using co-occurrence network analysis of the core microbiome, we identified the pathobiome comprised of the pathogen and key associate phylotypes in the cotton microbiome. Isolation and application of some negatively correlated pathobiome members provided protection against plant infection. Importantly, our field survey from multiple cotton fields validated the pattern and responses of core microbiomes under FOV infection. This study advances key understanding of core microbiome responses and existence of plant pathobiomes, which provides a novel framework to better manage plant diseases in agriculture and natural settings.

## Introduction

Plant diseases caused by a diverse range of soil-borne pathogens are major constraints of primary productivity worldwide (Strange and Scott, 2005). A recent study predicted that climate change would significantly increase the abundance of potential soil-borne pathogens in global soils with potential negative impacts on food security (Delgado-Baquerizo *et al*., 2020; Chaloner *et al*., 2021). Typical methods of disease control in agriculture rely heavily on synthetic chemicals (e.g., biocides), cultivar selection, plant breeding, and management practices such as crop rotations (Yang *et al*., 2008; Ulloa *et al*., 2020). However, for several soil-borne fungal pathogens, the use of chemical and plant breeding control measures is becoming increasingly ineffective (Hollomon, 2015; Lucas *et al*., 2015). Additionally, management approaches (e.g., crop rotation) can have significant economic consequences, particularly where crop selection is limited by environmental conditions and where pathogens can survive for long-periods of time in soils in the absence of preferred hosts (Göre *et al*., 2009). Furthermore, in field conditions, disease incidence is a function of the interactions between the host, pathogen, and environmental conditions including plant/soil associated microbiota (Trivedi *et al*., 2017a; Trivedi *et al*., 2017b). Therefore, deepening our understanding of how pathogens survive, respond to management practices, and interact with other soil biota, is considered an important priority. Such knowledge is needed to develop effective and environmentally sustainable approaches to manage plant diseases (Padda *et al*., 2017; Qiu *et al*., 2019).

In recent years, the role of indigenous soil microbiomes in suppressing plant diseases in both glasshouse and field conditions have been widely reported (Schlatter *et al*., 2017; Bakker *et al*., 2018; Berendsen *et al*., 2018; Kwak *et al*., 2018; Carrión *et al*., 2019; Wei *et al*., 2019; Liu *et al*., 2020). Given their intimate contact with the plant environment, rhizosphere microbiomes (i.e. soil associated directly with plant roots) are proposed to play an even more important role in protection against pathogen invasion than bulk soil microbiomes (Singh *et al*., 2020; Trivedi *et al*., 2020). Rhizosphere microbiomes can act to protect the plant against soil-borne pathogens via several mechanisms, including direct competition with pathogens for resources and space, active production of antibiotics effective against the pathogen, or indirectly by priming the host’s immune system (Leoni *et al*., 2020; Liu *et al*., 2020; Trivedi *et al*., 2020). For example, recent work reported an enrichment of *Stenotrophomonas rhizophila* (SR80) in the wheat rhizosphere and root endosphere in response to *Fusarium pseudograminearum* infection. Such enrichment contributed to plant disease tolerance via interacting with plant defence signalling pathways (Liu *et al*., 2021a). Similarly, plant cultivar dependent pathogen suppression (e.g., tomato) was reported to be linked with unexpected selection of specific bacterial species in the root environment leading to disease suppression (Kwak *et al*., 2018). In addition to specific microbial species being associated with increased resistance to plant pathogens, microbial diversity, and community structure have also been linked with disease suppression (Hu *et al*., 2016; Trivedi *et al*., 2020). It is likely that plant microbiomes utilise multiple biotic mechanisms to defend against pathogens which may vary between different pathogens, management practices and climatic conditions. Such complexity makes it challenging to identify and generalise the direct role that microbial communities play in disease suppression. Therefore, unravelling the role of microbial diversity and composition in disease protection, as well as their interaction with plant hosts and pathogens are crucial in the context of better understanding of disease incidence and its management.

The plant rhizosphere microbiome assembly is dynamic and therefore identifying the persistent (core) members of the plant microbiome has been proposed as a tool to advance understanding of soil-plant-microbiome interactions (Benson *et al*., 2010; Ainsworth *et al*., 2015; Chen *et al*., 2018; Schlatter *et al*., 2020) and key host functions (both negative and positive) provided by rhizosphere microbiomes (Shade & Handelsman, 2012; Singh *et al*., 2020). The core microbiome is comprised of members of microbial assemblages commonly present in hosts or within particular niches of a broad host community (Turnbaugh *et al*., 2007). In the context of the rhizosphere, the core microbiome of a plant species represent persistent microbial taxa associated with their host rhizosphere soils, regardless of their environmental conditions, geographical locations or management practices (Lemanceau *et al*., 2017). It has been reported that beneficial members of the core microbiome are critically involved in plant performance (Chen *et al*., 2018), and therefore, any direct and indirect changes in the core microbiome induced by disease can potentially alter host functions and phenotypes related to disease resistance and/or growth (Xu *et al*., 2018; Kaushal *et al*., 2020). Furthermore, some members of the core microbiome may provide an effective source of biocontrol agents against plant disease because of their ability to rapidly colonise plant surfaces and inner tissues and act as a first line of defence against invading pathogens; however, some members can also promote invasion of plant pathogens (Qiu *et al*., 2019; Singh *et al*., 2020). Therefore, the identity of the core members of the rhizosphere soil microbiome, their function, and response to biotic and abiotic drivers, is critically important to advance our understanding of the role of the soil microbiome in host performance.

The pathobiome, which is an emerging concept, considers disease as the manifestation of multiple interactions among several microbial species, including pathogens, which affect the health and disease status of the host (Vayssier-Taussat *et al*., 2014; Sweet & Bulling, 2017; Bass *et al*., 2019). Specifically, co-occurrence network analysis has been used previously to understand microbial interactions at interkingdom scales and to decode the influence of pathogen invasion on the indigenous microbial communities (Erlacher *et al*., 2014; Wei *et al*., 2015; Bass *et al*., 2019; Pauvert *et al*., 2020). Given that the pathobiome can be dynamic under different host and environmental conditions, using core members of the rhizosphere soil microbiome to identify the pathobiome could pinpoint microbial species that interact consistently with the target pathogen. However, both the existence of the pathobiome and its role in plant disease occurrence are yet to be fully explored.

The aim of this study was to identify the response of plant rhizosphere and bulk soil microbiomes to pathogen invasion and to determine whether application of biocontrol agents can resist the impact of pathogens. Furthermore, we aimed to identify the existence of pathobiomes in plant diseases. We selected cotton as our model plant host because globally it is a major cash crop, but is subject to significant loss of productivity globally (Davis *et al*., 1996; Davis *et al*., 2006) as a result of destructive disease caused by the fungal pathogen*Fusarium oxysporum* f. sp. *vasinfectum* (thereafter FOV). We combined glasshouse and field studies to first analyse the soil and rhizosphere microbiome of cotton plants (*Gossypium hirsutum*) in the presence and absence of FOV. We then examined the impact of biocontrol agents on soil and rhizosphere microbiomes. Using co-occurrence network analysis of the core microbiome, we identified the pathobiome. Importantly, findings from our glasshouse work were validated using extensive field surveys, where microbiome and pathogen data were collected from multiple cotton fields that were infected with FOV or were FOV-free.

## Results

### Effect of pathogen and biocontrol agents on plant performance and soil and rhizosphere microbiomes

To test the impact of FOV on cotton plant performance and associated plant microbiomes, and to examine the protective effect of biocontrol agents, three treatments were established in the glasshouse experiment: FOV inoculated (F); FOV plus biocontrol treatment (FB); and a non-treated control (C). Four widely used cotton cultivars (V1 = CIM 448, V2 = Siokra L23, V3 = CS 50, V4 = DP 16) and two soil types (clay soil and clay-sandy soil, detailed information described below) were used with three replicates for each treatment combination. Overall, we found the treatment using the biocontrol agent reduced the level of FOV infection. In F treatment, 25% of plants were diseased as confirmed by the observation of brown rings in the cross-sections of stems, while application of synthetic biocontrol community reduced infection to 13.3% in FB treatment (Figure S1). The quantitative PCR (qPCR) confirmed a significantly higher pathogen abundance in FOV-inoculated soil than uninoculated soil across all samples at seedling stage (P < 0.05, Figure S2). Disease incidence was found higher in in the presence of pathogen (Figure S1), while no significant effect of pathogen treatment on seed germination rate and plant height varied across different cotton cultivars (P > 0.05). Cotton productivity was slightly lower (~3 %) in the presence of the pathogen (Figure S1), but differences were not statistically significant (P > 0.05). In cotton soil microbial communities, samples were rarefied to equal sequencing depth per sample, resulting in a total number of 4,271,083 bacterial sequences spanning 13,445 OTUs and 13,650,185 fungal sequences spanning 7,031 OTUs. Although rarefaction curves of the bacterial sequences did not reach plateau, an average of 11,129 reads per sample were used, with approximately nine times of average richness per sample, suggesting good coverage of the microbial diversity present (Figure S3). Despite the significant differences between the two soil types and between bulk and rhizosphere soils (Figure S4A & B), we mainly focused on the variation of microbiomes under different treatments to evaluate the effect of pathogens and biocontrol agents.

In the glasshouse experiment, no significant difference was found in microbial diversity across different treatments in bulk soil samples (P > 0.05, Table S4, Figure S5A & B). In the rhizosphere soils, bacterial communities had significantly higher Shannon and Simpson indices in F treatment compared to bacterial communities in C and FB treatments (P < 0.05, Table S4, Figure S5A). In combination with qPCR results, the presence of FOV in bulk soil may increase Shannon and Simpson indices (i.e. increase community evenness). For fungal communities, in both rhizosphere and bulk soils, a lower Chao1 index was found in the FB treatment compared to C and F treatments (P < 0.05, Figure S5B, Table S4), implying that application of biocontrol agents might reduce the fungal richness. There were no significant differences in either bacterial or fungal alpha diversity indices observed in field samples (P > 0.05, Table S4, Figure S5C & D).

Furthermore, there was no significant treatment effect, or of their interaction on either bacterial or fungal communities associated with bulk soil (P > 0.05, Table S5). However, there were significant differences in the interaction between treatments, soil type and time in the rhizosphere bacterial communities and the interaction between soil and time in the rhizosphere fungal communities (P < 0.05, Table S5). Specifically, a consistent pattern was found across the two soil types at seedling stage where the rhizosphere bacterial community in treatments C and FB were different from treatment F, but no significant difference was found between C and FB (P < 0.05, Table 1A). This pattern was also aligned with the alpha diversity results (Table S4). Interestingly, no difference was observed between treatments in either bacterial or fungal communities in rhizosphere samples at flowering stage (P > 0.05, Table 1A). Principal coordinates analysis (PCoA) also demonstrated that rhizosphere bacterial communities were mainly differentiated by treatment at the seedling stage, but tended to be similar between treatments at the flowering stage (Figure 1).

**Figure 1.**
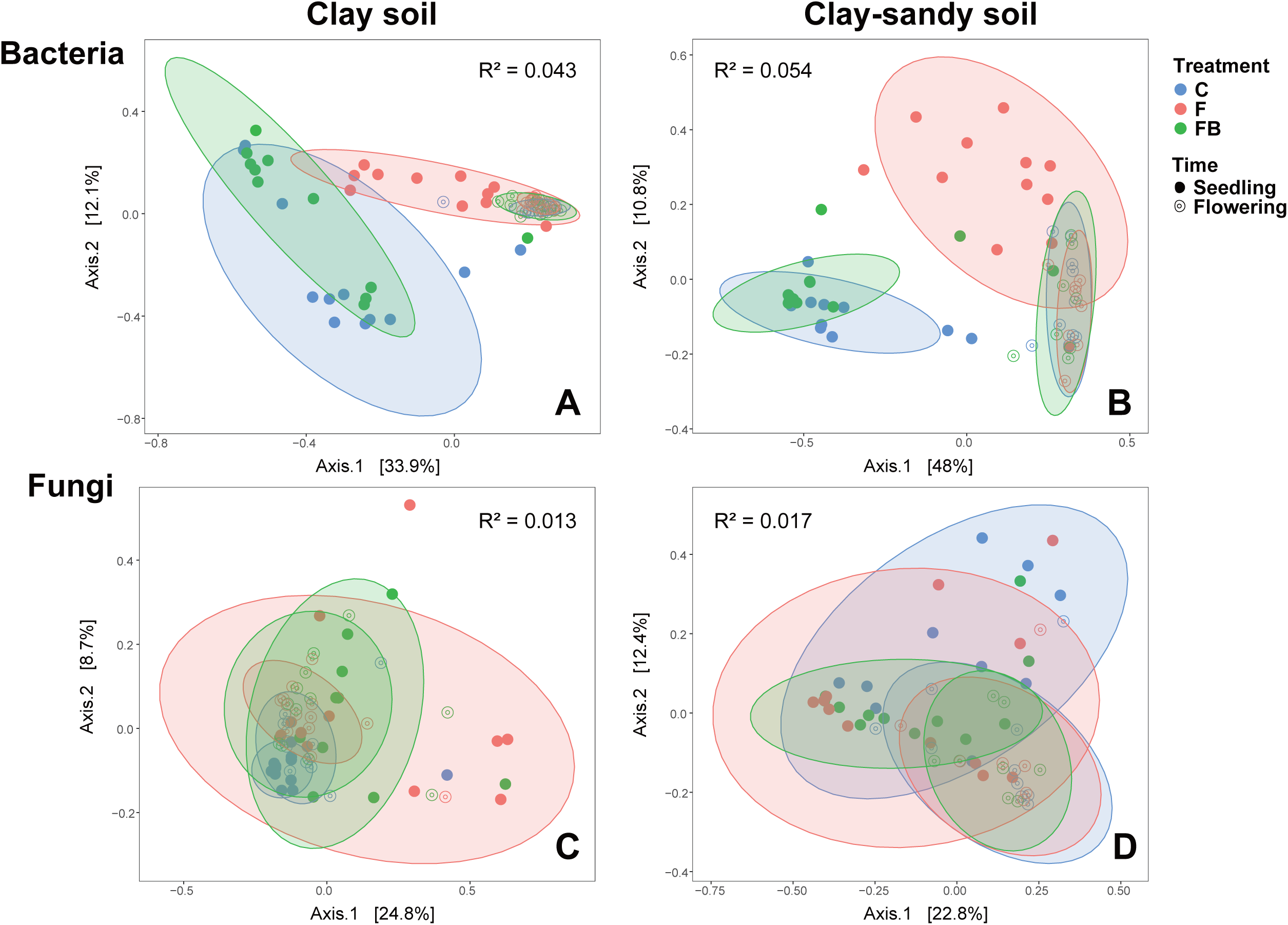
Principal Coordinates Analysis (PCoA) plot using Bray-Curtis distance matrix on bacterial (A & B) and fungal (C & D) communities in different soil types (A & C: clay soil, B & D: clay-sandy soil). C = control treatment (light blue), F = FOV treatment (light red), FB = FOV + biocontrol treatment (light green). Solid circles indicate samples at seedling stage, and open circles indicate samples at flowing stage. Ellipses represent 95% confidence interval.

Data from the field samples showed consistent and significant differences in both bacterial and fungal communities of diseased plants in diseased fields compared to plants in healthy fields in bulk soil samples from the Macquarie region (P < 0.05, Table 1B, Table S6), while rhizosphere communities of all plants (symptomatic and asymptomatic) from diseased fields were different from healthy fields in St George region (P < 0.05, Table 1B, Table S6). Principal coordinates analysis (PCoA) plots showed that location, along with the health status of plants, directly influenced the variation of microbial communities, but no differences were evident between symptomatic and asymptomatic plants from within disease fields (Figure S4C & D).

### Response of microbial phylotypes to diseases incidence

The use of the biocontrol significantly altered the impact that the pathogen had on rhizosphere microbiomes. Results showed that bacterial communities differed significantly between C and F, FB and F, but not between FB and C during seedling stage (Table 1). To further investigate the treatment effect on representative OTUs in the rhizosphere microbiome at the seedling stage, microbial communities were analysed at the OTU level using LEfSe. Overall, distinct rhizosphere bacterial OTU indicators were found between treatments irrespective of growth stage or soil type (Figure 2A). Specifically, indicator OTUs representative of treatment F mainly belonged to Actinobacteria and Bacteroidetes phyla, and to the class Alphaproteobacteria. Representative OTUs of C and FB treatments belonged to the phylum Firmicutes and the class Gamma proteobacteria, respectively (Figure 2A). In field samples, OTU indicators of healthy and diseased fields belonged mainly to the phylum Proteobacteria and phylum Firmicutes, respectively (Figure 2B). Despite contrasting microbial compositions between glasshouse and field rhizosphere soils, similar trends of enrichment of OTUs belonging to Proteobacteria in treatment FB and healthy fields were observed (Figure 2A & B). For fungal indicators in the glasshouse experiment, a majority of representative OTUs of treatment F belonged to the phylum Zygomycota, whereas OTUs of treatment FB were found in Ascomycota and Basidiomycota phyla (Figure 2C). In field samples, although indicator OTUs from different phenotypes of cotton plants in diseased field were mainly found from the phylum Ascomycota, a majority of representative OTUs from healthy plants belonged to Onygenales, Hypocreales and Microascales orders, while OTUs from diseased plants belonged to Xylariales and Eurotiales orders (Figure 2D).

**Figure 2.**
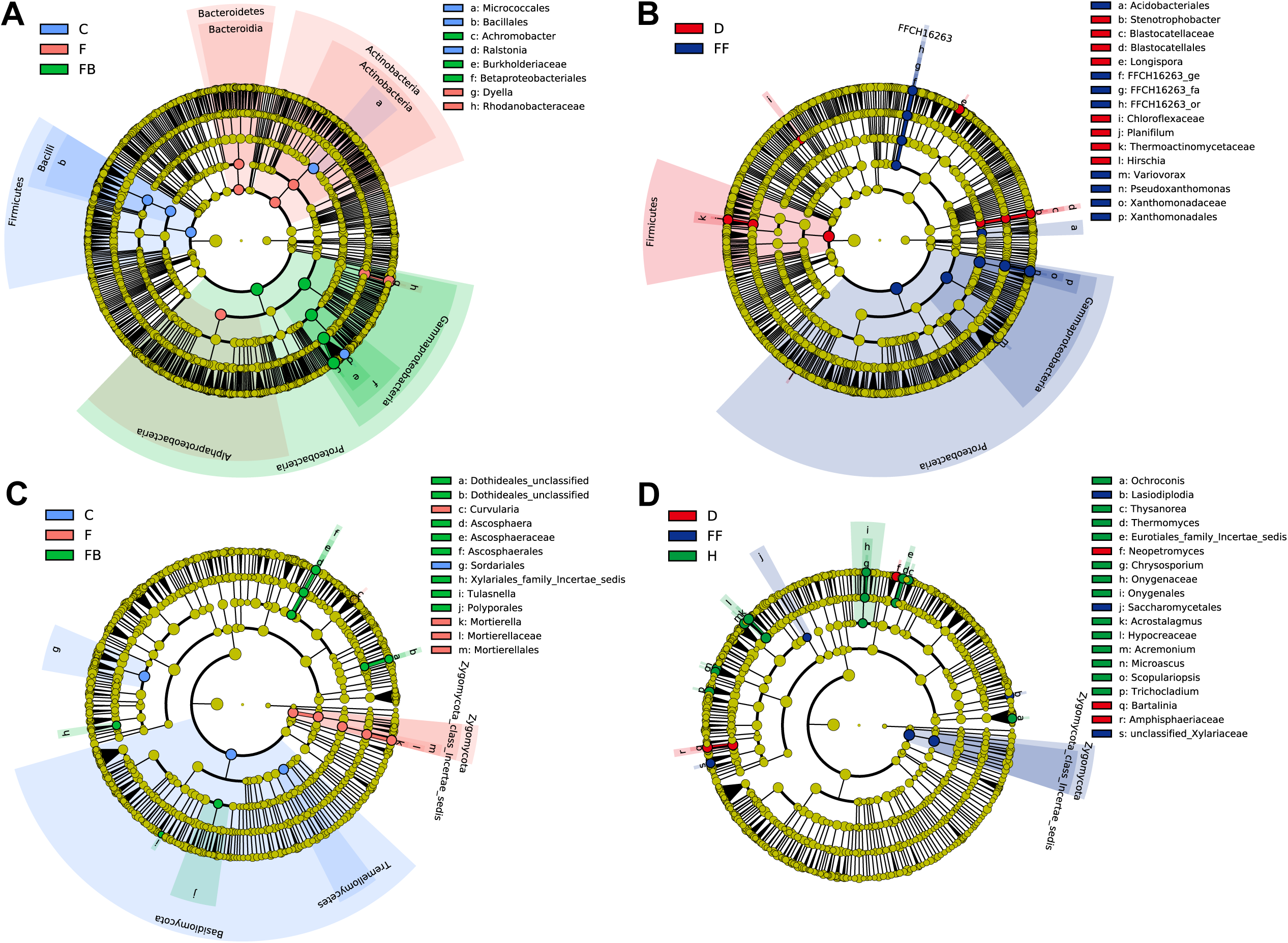
Cladogram of bacterial (A & B) and fungal (C & D) communities based on LEfSe analysis (LDA effect size cutoff = 3.0) in glasshouse (A & C) and field (B & D) samples. Microbial markers at different taxonomic levels are highlighted with colours based on treatments or plant phenotypes: C = control treatment (light blue), F = FOV treatment (light red), FB = FOV + biocontrol treatment (light green), D = diseased plants in diseased field (red), H = healthy plants in diseased field (green), FF = healthy plants in FOV-free field (navy). Notably, there was no bacterial indicator found (LDA effect size ≥ 3.0) in H group in field samples (B).

### Disruptions of cotton core members of rhizosphere microbiome under FOV attack

To identify the core members of the rhizosphere microbiome, OTUs that were present in >75% of samples of a particular treatment group were extracted as the core microbiome. When OTUs from all available glasshouse treatments were combined, only 72 (0.98%) of total bacterial, and, 53 (1.15%) of total fungal OTUs from all treatments were considered as core OTUs, but represented 56.79% and 4.36% of total relative abundance, respectively (Figure 3A). In field samples, 90 (1.39%) of total bacterial and 96 (2.48%) of total fungal OTUs across all samples were identified as core, accounting for 51.55% and 2.96% of total relative abundance, respectively (Figure 3). Given inconsistency and lack of dominance of core fungal biota, we mainly focus on bacterial core microbiomes for further analyses. There was significant overlap between field and glasshouse data. Thirty-three core OTUs were shared between glasshouse and field samples, which were mainly members of the phyla Actinobacteria, Firmicutes, Gemmatimonadetes, Proteobacteria and Ascomycota (Figure 4, Table S7), suggesting a similar pattern of core taxa could be found in cotton rhizosphere irrespective of the geographical location, soil type or plant vegetation stage. Relative abundance of bacterial core OTUs showed that dominant OTUs (e.g., OTU00001, OTU00002 and OTU00007) decreased significantly in glasshouse core rhizosphere microbiome under FOV treatments. This negative impact of FOV on relative abundances of these OTUs was protected by biocontrol treatment (Figure 3A). Similarly, dominant rhizosphere core OTUs were higher in healthy plants, particularly healthy individuals in diseased fields (e.g., OTU00002, OTU00028 and OTU00036, Figure 3B). Importantly, OTU00002 (*Bacillus* sp.) was consistently found in both glasshouse and field samples, suggesting this bacterial species could be closely related to plant fitness under *Fusarium* invasion. In fungal communities, OTUs shared between glasshouse and field samples were mainly from genera *Fusarium, Acremonium, Chrysosporium* and family *Chaetomiaceae* (Table S7). Full information of core OTUs and relative abundances are listed in Table S7 and S8.

**Figure 3.**
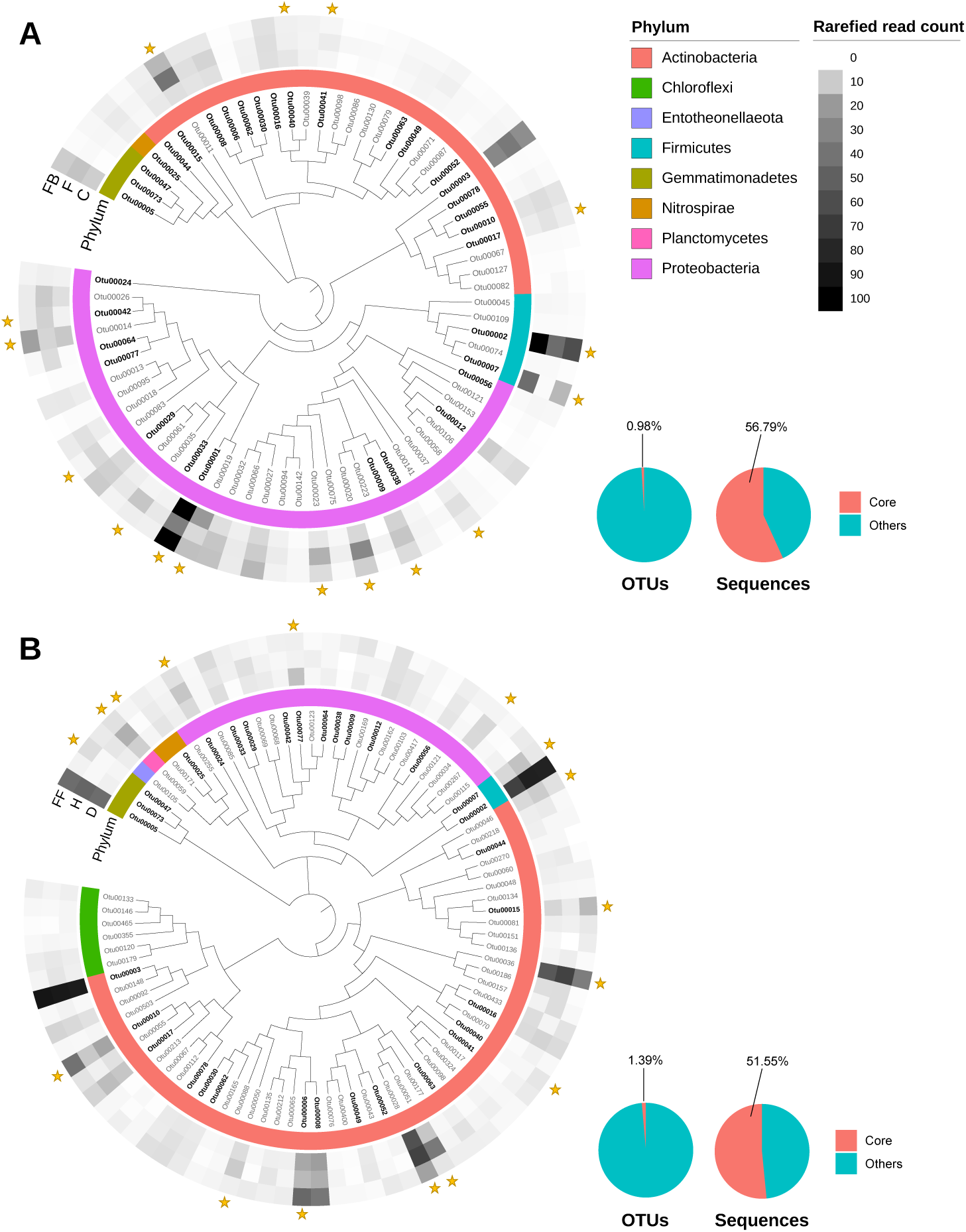
Bacterial core OTUs in glasshouse (A) and field (B) samples. Phylogenetic trees were constructed using maximum likelihood method based on 16S rRNA gene V5-V7 region (799F-1193R). The outer strips indicate relative abundances of each OTU under different groups, and the inner coloured strip indicate bacterial phyla of the OTUs.

**Figure 4.**
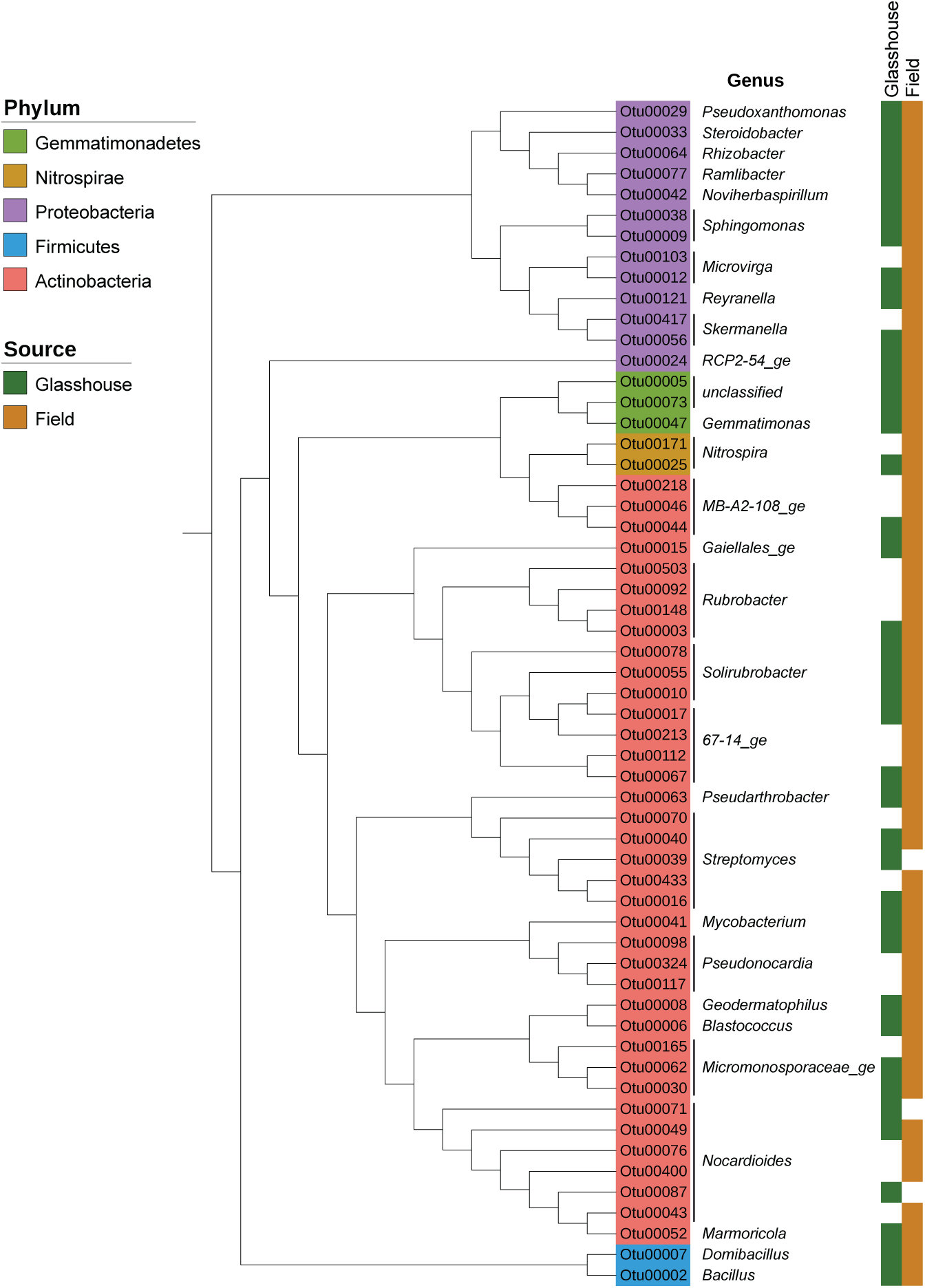
Universal core bacterial genera with inclusive OTUs present across both glasshouse and field samples. Phylogenetic trees were constructed using maximum likelihood method based on 16S rRNA gene V5-V7 region (799F-1193R). Core OTUs presented in glasshouse samples are labelled with green blocks, and core OTUs presented in field samples were labelled with brown blocks.

### FOV pathobiome in fungal-bacterial networks

The co-occurrence network analysis of bacterial/fungal OTUs with strong correlations (| r | > 0.6) showed that fewer associations were found in the presence of biocontrol agent (FB, Figure 5A) compared to control (C) and FOV-treated (F) samples. Presence of FOV seemed to increase the bacterial positive associations in the glasshouse experiment compared to non-FOV treated soils (C vs F, Figure 5A, Table S9). We found similar results for healthy plants from healthy fields (FF, Figure 5B, Table S9). In addition, higher fungal-fungal associations were found in healthy plants, which was consistent in both glasshouse and field samples (Table S9).

**Figure 5.**
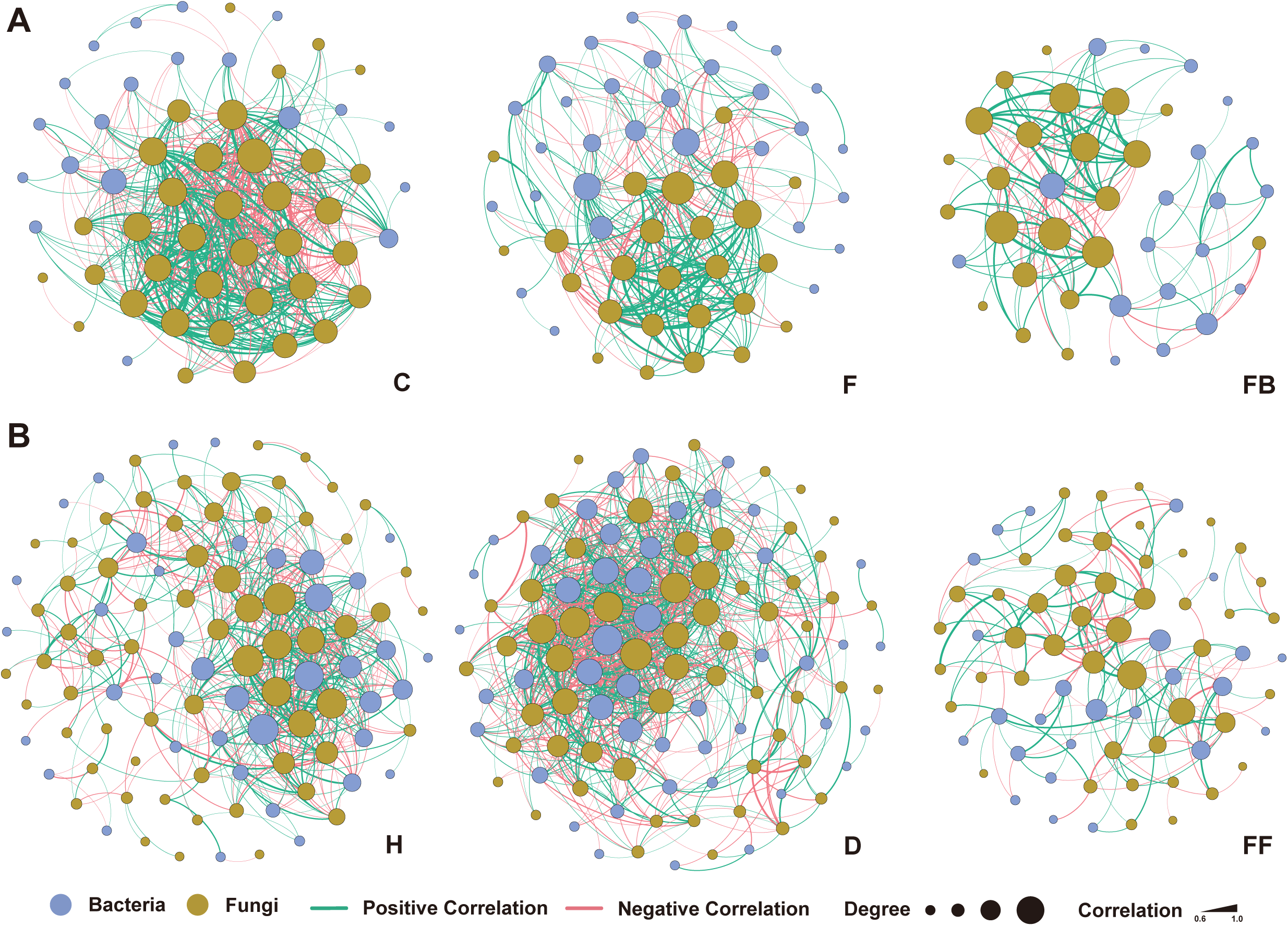
Co-occurrence network analysis of bacterial and fungal communities in glasshouse (A) and field (B) samples in different groups. Colours of nodes indicate bacterial (blue) and fungal (olive) OTUs, and colours of edges indicate positive (green) and negative (red) correlations. The size of nodes indicates the weight of the corresponding OTU (numbers of edges connected), and the weight of the edges indicates the degree of the correlation.

To identify the *F. oxysporum* pathobiome, OTUs associated with *F. oxysporum* were extracted from the network. In the glasshouse experiment, nine bacterial and 24 fungal OTUs were significantly (P < 0.05) associated and strongly correlated (| r | > 0.6) with *F. oxysporum* (Figure 6A), while 23 bacterial and 32 fungal OTUs were significantly (P < 0.05) associated, and strongly correlated (| r | > 0.6) with *F. oxysporum* (Figure 6B) in field samples, which identified as key pathobiome. Overall, an overlap in bacterial pathobiomes of glasshouse and field samples was observed from members of Actinobacteria, Firmicutes and Proteobacteria phyla. Similarly, the overlap in fungal pathobiomes observed at phylum level were Ascomycota and Zygomycota. A number of fungal OTUs were consistently found in the pathobiome of both glasshouse and field samples, including OTUs classified as *Acremonium alternatum, Chrysosporium pilosum*, sp., *Microascaceae* sp., *Eurotiomycetes* sp., and one unclassified Ascomycota that was negatively associated with *F. oxysporum* (Figure 6), which may be considered as potential antagonists against *F. oxysporum*. We successfully isolated five bacteria with antifungal activity against FOV that were identical to key members of pathobiome (based at > 97% similarity of sequences) identified by network analysis (Table S10), suggesting that the corresponding bacterial taxa in the pathobiome have a role in pathogenesis.

**Figure 6.**
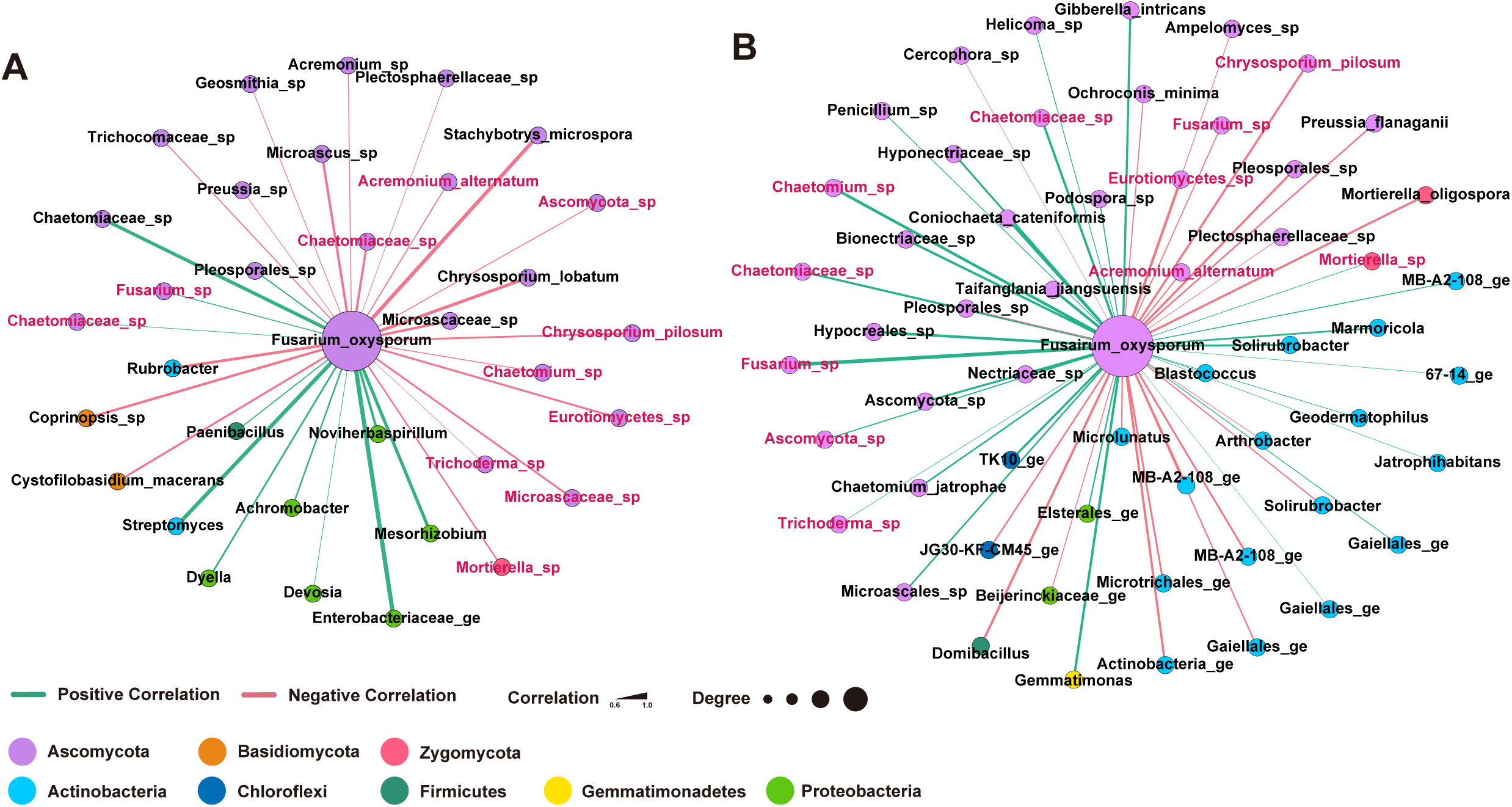
Pathobiome based on spearman correlations between *Fusarium oxysporum* and other microbial taxa in glasshouse (A) and field (B) samples. Nodes only with strong correlation (P < 0.01, | r | ≥ 0.6) are shown. Colours of nodes indicate the microbial phyla, and colours of edges indicate positive (green) and negative (red) correlations. Microbial OTUs consistently present in both glasshouse and field samples are highlighted with red.

## Discussion

### Fusarium infection shifts early microbial colonisation

Our results provide empirical evidence for the direct impact of pathogens on rhizosphere soil microbial communities at the seedling stage. Such shift of microbial communities influenced by fungal pathogens has be observed in other plants, such as banana, wheat and watermelon (Shen *et al*., 2018; Araujo *et al*., 2019; Wang *et al*., 2019). Biotic environmental stresses (e.g. plant diseases) can potentially trigger changes in rhizosphere microbial communities via direct interactions with invading pathogens or indirectly via altered plant root exudation and changes in plant physiology (Gu *et al*., 2016; Liu *et al*., 2020). The evidence that seedlings are more susceptible to FOV infection aligns with other plant pathogens (Develey-Rivière & Galiana, 2007), likely due to the lack of a fully developed plant immune system, which is both plant and microbial-mediated. In contrast, little difference in the rhizosphere microbiome was observed at the flowering stage, which indicates that plants are able to regulate rhizosphere communities via secondary metabolisms at later stages of development, irrespective of the status of disease incidence at seedling stage (Liu *et al*., 2020). Our glasshouse results were supported by field data, where asymptomatic plants in diseased fields had similar microbial assemblages to plants in healthy fields (Table 1B). Our finding is also consistent with the report that FOV has stronger impact on plants at the early stage of development (Bugbee, 1970), which means once plants escape the severe disease, they are able to maintain fitness and productivity at later stages. In bulk soils, differences of microbial communities were driven by the soil types and plant cultivars, indicating that cotton plants may be more responsible for the assemblage of the rhizosphere microbiome than bulk soil in the Macquarie fields (Table 1B, Hamonts *et al*., 2018; Xiong *et al*., 2021a; Xiong *et al*., 2021b). Markedly, in the glasshouse experiment, no significant difference was observed in rhizosphere microbial community between healthy and FOV with biocontrol treatments, implying that the biocontrol agent was able to protect plants from FOV attack. The effect of biocontrol agent could be further supported by a bacteria-only treatment, similar to a recent study on wheat (Liu *et al*., 2021b). Overall, our results demonstrated that FOV can introduce significant shifts in the rhizosphere microbiome at an early stage of plant development, when plants are most vulnerable to pathogen attack, but the use of a biocontrol can resist the negative impact of FOV.

### Persistent members of plant microbiome are responsive to FOV infection

This study provides the first report on the cotton rhizosphere core members of microbiome and its temporal and disease-induced changes in both glasshouse and field conditions. It consisted of a wide range of taxa from Actinobacteria, Firmicutes, Proteobacteria (mainly Alphaproteobacteria and Gammaproteobacteria), Ascomycota and a few taxa from other phyla. Although rhizosphere samples collected in the field were not treated identically as in glasshouse, which could potentially cause variations of microbial communities in the analysis, there were common bacterial genera observed as core microbiota such as *Bacillus, Burkholderia, Pseudomonas, Rhizobium, Streptomyces* and *Xanthomonas*, aligned with other crops such as sugarcane, canola and wheat (Hamonts *et al*., 2018; Lay *et al*., 2018; Schlatter *et al*., 2020). However, there were a few taxa exclusively found in cotton rhizosphere microbiomes, such as *Azotobacter, Microvirga, Skermanella*, and *Steroidobacter*. These findings indicate that core members of microbiomes can consist of both generalist (common in many plant species) and specialist (plant species specific) members, which may synergistically respond to and play crucial roles in plant interactions with pathogens (Trivedi *et al*., 2020).

Specific members of the rhizosphere core microbiome could possibly be altered under significant biotic stresses (Erlacher *et al*., 2014; Hamonts *et al*., 2018). Our results revealed that FOV-induced changes in core members of the rhizosphere microbiome can be observed in both identity and abundance, especially in members of Actinobacteria, Alphaproteobacteria, Gammaproteobacteria and in fungal taxon *Fusarium*. As the ITS region was selected for fungal community analysis, confirming whether the *Fusarium* OTUs are inoculated FOV or other *Fusarium* strains, including non-pathogenic *Fusarium*, would be difficult. Remarkably, relative abundance of a wide range of potential biocontrol agents were enriched in the rhizosphere under disease conditions, such as *Streptomyces* sp., *Microbispora* sp. and *Nocardioides* sp. in the phylum Actinobacteria (Misk & Franco, 2011; Palaniyandi *et al*., 2013; de Jesus Sousa & Olivares, 2016; Essarioui *et al*., 2017; Cabrera *et al*., 2020), and *Pseudomonas* sp. and *Sphingomonas* sp. in the class Gammaproteobacteria (Babu *et al*., 2000; Wachowska *et al*., 2013). These findings potentially support the recent evidence for the ‘cry for help’ strategy of plants under pathogen attacks (Liu *et al*., 2020), and may suggest that such a strategy of accumulating beneficial microbes in plant microbiomes could be common in many plant species.

Importantly, while differences in the rhizosphere microbiome between treatments were mainly observed at the seedling stage, we found that the relative abundance of core members of the rhizosphere microbiome also differed between control and FOV treatments, indicating that FOV fundamentally changed the core members of the cotton rhizosphere microbiome, regardless of the soil type and plant cultivar. This pattern was also found in our extensive field surveys, where flowering cotton plants were investigated and a consistent shift in certain core members of the rhizosphere microbiome was observed from plants from diseased and healthy fields (e.g. *Bacillus* sp., *Solirubrobacter* sp., *Streptomyces* sp., etc.). Previous studies have demonstrated that a shift in core members of the microbiome and their functions can have prominent negative impacts on plant fitness, although the magnitude of the impact likely varies depending on plant species and microbial taxa (Vandenkoornhuyse *et al*., 2015; Chen *et al*., 2018; Hamonts *et al*., 2018). The core identity of microbiomes is emerging, so a better understanding of the mechanisms that drive shifts in core microbiomes, and consequences for host functions, should be the focus of future research. Such knowledge will foster pathways for harnessing plant microbiomes to improve plant health and productivity in both natural and agricultural settings.

### Identity of pathobiome

Members of a microbiome continuously interact with each other and maintain underground microbial ecosystems, providing feedbacks to plant productivity and soil health (Kulmatiski & Beard, 2011; Lamb *et al*., 2011). The microbial network analyses demonstrated more correlations between microbes in FOV-infected than in non-infected samples of both glasshouse and field data. While the correlation patterns in the field samples paralleled with the glasshouse samples, microbial networks were altered in FOV-infected samples, which is consistent with previous findings for other pathogens such as soil-borne *Ralstonia solanacearum* (Rybakova *et al*., 2017; Wei *et al*., 2018). Hence, the microbiome network status in the rhizosphere could influence plant fitness and be used to predict threats from potential pathogens.

The “pathobiome” concept has expanded our view from a single microorganism as a disease agent to a broader perspective of communities that co-affect a particular disease (Vayssier-Taussat *et al*., 2014; Sweet & Bulling, 2017), leading to a series of innovative pathobiome studies in human (Krezalek *et al*., 2016), livestock (Tufts *et al*., 2020), large marine organisms (Sweet *et al*., 2019) and some plants (Doonan *et al*., 2019). Our study has used the core members of microbiome to build the pathobiome of *F. oxysporum*, by initially narrowing the entire microbial community to the ubiquitous single species before identifying the pathobiome. This approach accommodates the identification of a core network, which can avoid screening the rare/ transitional taxa in the environment that occasionally appeared in soil and plant samples (Thomas *et al*., 2016). While positive associations between pathogens and other microbiota may help pathogens to cause disease (Hoffman & Arnold, 2010; Jakuschkin *et al*., 2016), they may also represent a common response in the stressed plants under pathogen attack (Sweet *et al*., 2019). On the other hand, negative associations can be considered as potential biocontrol agents against specific pathogens (Pauvert *et al*., 2020). Following the network analysis approach (Pauvert *et al*., 2020), we extracted 78 bacterial and fungal OTUs from core network as the putative pathobiome of *F. oxysporum*, of which 32 bacterial taxa were identified as key members of the pathobiome, including 19 taxa from the phylum Actinobacteria, suggesting these taxa potentially provide antifungal activities at early stages of the FOV infection (de Jesus Sousa & Olivares, 2016; Goudjal *et al*., 2016). We were able to isolate five members of the pathobiome and they all showed antifungal activities against FOV suggesting these microbes have a direct role in pathogenesis and that negatively associated microbial taxa of the pathobiome can be considered as candidate biocontrol agents relevant to disease suppression. By identifying and isolating more members of the pathobiome (which either help or hinder pathogenesis), targeted disease management, including biocontrol agents against pathogens, could potentially be developed. However, further mechanistic understanding is required to apply such tools in agriculture and forestry settings. For example, a better understanding is needed regarding the mechanisms by which microbial taxa support or supress disease development, and the ability to discriminate between taxa generating the effect (e.g., disease suppresser) from those present as opportunists.

### Biocontrol agents resist the impact of FOV on the rhizosphere microbiome

There were significant differences between the rhizosphere microbiomes of control and FOV treated samples in the glasshouse experiment. This result was also consistent with our field data, which showed differences in plant rhizosphere microbiomes of healthy and diseased fields, providing evidence that the change in microbiome was driven by FOV infection (Saravanakumar *et al*., 2017; Wei *et al*., 2018). However, FOV treated samples in the presence of biocontrol agents prevented such infection-induced changes in the rhizosphere in the glasshouse experiment. The variation of microbial communities between biocontrol treated and untreated rhizosphere samples under pathogen attack in our studies are consistent with previous work on wheat (Araujo *et al*., 2019) and tomato (Elsayed *et al*., 2020), suggesting that biocontrol agents can effectively protect the rhizosphere microbiome from significant impacts by pathogens.

Microbial interactions are crucial for disease development. Previous studies have provided evidence that negative interactions between pathogen and plant associated microbes can significantly enhance a plants ability to defend itself from the pathogen invasion (Gajbhiye *et al*., 2010; Goudjal *et al*., 2016; Araujo *et al*., 2019). We found that apart from the bacterial taxa in the phylum Actinobacteria, additional bacterial taxa with antifungal properties were also abundant in the FB treatment, such as *Bacillus* sp., and *Brevibacillus* sp. from the phylum Firmicutes (Khan *et al*., 2017; Jangir *et al*., 2018; Johnson *et al*., 2020), and *Pseudomonas* sp. from the class Gammaproteobacteria (Babu *et al*., 2000; Arya *et al*., 2018). These taxa have been commonly used as potential biocontrol agents (Szczech & Shoda, 2006; Chen *et al*., 2017; Sun *et al*., 2017), and were frequently detected in our study, suggesting that the potential bacterial-fungal interactions disrupted by FOV were largely restored by the biocontrol treatment.

In the core members of the microbiome and network analyses presented here, FB treatment had less core OTUs / associations relative to control and FOV treatments, but most of which overlapped with the control treatment. This result is also supported by the beta diversity analysis (C ≠ F, FB ≠ F, C = FB), which showed that core members of microbiome and network associations in FB remained similar to control, and, shifts in microbial community structure were minimised at the seedling stage. Network analysis does not identify positive and negative interactions, but rather provides associations. Current literature suggests that both interpretations are possible (Jakuschkin *et al*., 2016; Sweet *et al*., 2019) and manipulative experiments would be necessary in order to identify whether these associations were positive or negative interactions. Nevertheless, our work provides critical empirical evidence of microbial responses of disease incidence and provides a conceptual framework to develop and test targeted research on plant-microbial interactions under biotic stresses.

Overall, we provide evidence for the existence of a core microbiome of cotton which contains both generalists (common in many plant species) and specialists (plant species specific) members. We demonstrated that pathogen invasion results in a consistent shift in the rhizosphere microbiome and network associations, both in glasshouse and field conditions. FOV-induced shifts in the host and microbiome were prevented by treatment with biocontrol agents. Furthermore, we provide evidence for the existence of a pathobiome in plants, which provides a tool to identify microbiota that are positively associated with and facilitate pathogenesis. Similarly, negatively associated microbial taxa of the pathobiome may represent potential biocontrol agents. This opens new avenues to understand plant-microbial and microbial-microbial interactions in pathogenesis and to develop new approaches for sustainable management of plant diseases.

## Experimental Procedure

### Glasshouse Experiment

#### Construction of synthetic community with biocontrol properties against FOV

A glasshouse experiment was performed to test the impact of FOV on cotton plant performance and associated plant microbiomes, and to examine if the use of biocontrol agents could resist pathogen impacts on the plant microbiome and pathobiome. We established three treatments: FOV inoculated (F); FOV plus biocontrol treatment (FB); and a non-treated control (C). Four widely used cotton cultivars (V1 = CIM 448, V2 = Siokra L23, V3 = CS 50, V4 = DP 16) and two soil types (clay soil and clay-sandy soil, detailed information described below) were used in the design with three replicates for each treatment combination.

Bacterial biocontrol and FOV isolates were obtained from our laboratory culture collections. Bacterial strains were previously isolated from the rhizosphere collected from a cotton field in Narrabri, NSW Australia, and FOV was sourced from our lab collection which was originally isolated from FOV-infected cotton. FOV was first revived on LB agar at 28 °C for 24 h, before subculturing on new agar plates for antagonistic assay with biocontrol agents. Taxonomic identity of the seven selected bacterial strains (Table S1) of the synthetic community, and FOV were characterised by 16S rRNA genes and ribosomal internal transcribed spacer (ITS) region using primer 27F/1492R (Lane, 1991) and ITS1/ITS2 (Ihrmark *et al*., 2012) by Sanger sequencing at Western Sydney Genomic Facility. Details of the biocontrol selection were described in the Supplementary File.

#### Seeds, biocontrol agents and seed treatment

Cotton seeds were first washed with 0.1% HgCl_2_ solution in a 50 ml falcon tube to remove the attached fungi from seed surface, and then washed with distilled water three times to remove the remaining HgCl_2_ (Ramakrishna *et al*., 1991). Seeds were then washed with 70% ethanol to remove attached bacteria, followed by washing three times in sterile water to remove residual ethanol. Each of the washing steps was conducted by hand shaking in 50 ml falcon tubes for 1 min.

Prior to planting, cotton seeds were treated with either sterile distilled water (C and F) or the consortium of the selected bacterial isolates (FB). For constructing the biocontrol synthetic community, bacteria were first grown together on an agar plate to test their compatibility. Selected bacterial strains were then sub-cultured in 100 ml LB broth to exponential growth phase by estimating the cell density with NanoDrop™ spectrophotometer at OD_600_ (OD_600_ at 0.5 to 0.6, NanoDrop 2000, Thermal Scientific™). Bacterial cultures were collected by centrifugation (4000 g, 10 min) and resuspended with sterile water to a final concentration of approximately 4 ×10^7^ CFU/ml. All suspended bacterial strains were mixed at equal concentration to produce the bacterial synthetic community used in the biocontrol treatments. Sterilised, dried cotton seeds were immersed either in bacterial synthetic community solution or distilled water for 30 min on a shaker (50 rpm) at room temperature.

Two types of soils were used in this study to provide variation in soil abiotic and biotic properties. The clay soil was collected from a cotton cultivation farm near Griffith NSW, Australia (34.16 S, 146.01 E) and the clay-sandy soil (thereafter sandy soil) was collected from cotton growing area at Western Sydney University farm facility, Richmond NSW, Australia (33.61 S, 150.74 E). The initial soil physiochemical parameters including nitrate, nitrite, pH, and soil moisture were measured and described in Table S2. Experimental pots (25 cm diameter by 30 cm depth) were filled with 3 kg soil. For the establishment of FOV treatment, FOV was sub-cultured in 200 ml full strength potato dextrose broth for 24 hours. The conidia produced were quantified with a haemocytometer under a microscope by averaging five times of counting. FOV conidia were centrifuged at 4000 g for 10 min, resuspended with sterile water and inoculated into each pot of soil at a final concentration of approximately 10^7^ CFU per gram of soil before being thoroughly mixed. For uninoculated control treatments, an equal amount of sterile water was added to the soil.

#### Experimental conditions and plant performance

Five seeds treated with biocontrol agent were sown in a FOV-inoculated pots, while untreated seeds were sown in both FOV-inoculated / uninoculated pots. Pots were randomly distributed in a naturally sun-lit, ambient [CO_2_], glasshouse bay. The temperature was set at 32 °C daytime, 28 °C morning/evening for 2 h, respectively, and 25 °C night-time, which are typical temperatures of cotton cropping regions in eastern Australia (http://www.bom.gov.au/). Sterilised water was supplied daily to maintain soil moisture above 70% Water holding capacity (WHC) and fertilisers (Thrive All Purpose Soluble Fertiliser, Yates) was supplied fortnightly following manufacturer’s instructions (see details in Table S2). Germination data were collected daily for 14 days, plant height data were collected weekly, and cotton productivity (lint weight) was recorded at final harvest (20 weeks post-sowing). Disease incidence was confirmed by the leaf wilting levels (healthy, light and severe) and the presence of brown rings in the cross section (Figure S7) by cutting off cotton stems 10 cm above ground using a stem pruner.

#### Quantitative PCR analyses of FOV abundance in bulk soil and the rhizosphere

To compare FOV load between samples, qPCR was carried out using FOV-specific primer pairs Fov-1 (5’-TGTAGGGGTTGTGGGTTTTTTTC-3’) and Fov-2 (5’-CCAACACACAACCGCACACGA-3’), which amplifies a 125-bp DNA fragment (Zambounis *et al*., 2007). Total fungi load in soils was quantified using the primer pairs ITS1f (5’-TCCGTAGGTGAACCTGCGG-3’) and 5.8S (5’-CGCTGCGTTCTTCATCG-3’) (Fierer *et al*., 2005). Reactions were performed in duplicate in a LightCycler 480 (Roche) using 5□µL 2X LightCycler 480 SYBR Green I Master mix, 15 pmol primer mix and 1µL template DNA (2.5□ng) in a 10 µL reaction volume. Thermal conditions for both genes consisted of 5 min at 95□°C for initial denaturation, 40 cycles of 15□s at 95□°C, 20 s at 61°C (FOV)/53°C (Fungi) for annealing and 30□s at 72□°C. FOV abundance in soils were estimated with the formula:

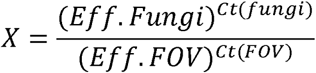

where Eff. (Fungi/FOV) is amplification efficiency for qPCRs, calculated by LinRegPCR 7.5 using raw amplification data for each sample (Ramakers *et al*., 2003). The Cts (Fungi and FOV) are threshold cycles, and X represents the percentage of FOV copy numbers existing in a sample.

#### Rhizosphere and bulk soil sample collection for the glasshouse experiment

For assessing soil microbiome changes by the pathogen infection, bulk and rhizosphere soil samples were collected at two time points: seedling (approximately two weeks after germination) and flowering (when more than half of the plants had flowers) stages. Briefly, approximately 2.5 g of bulk soil from topsoil (0-10 cm) of each pot was collected within a 10 cm radius from the plant stem using a sterilised spatula, and transferred into a 2 ml cryotube. For rhizosphere soil, one cotton plant from each pot was carefully uprooted from the soil and gently shaken to remove the excess attached soil, then submerged with TE buffer (10mM Tris-HCl, 1mM EDTA, pH 8.0) in a 15 ml falcon tube, gently shaking by hand for 30 s to wash off the rhizosphere soil. Washed plant roots were then transferred into another clean 15 mL falcon tube. The rhizosphere soil was collected by centrifugation at 4000 g for 10 min, with supernatant discarded. All collected samples were immediately stored at −80 °C until being processed for DNA extraction.

#### Field sampling

To further investigate microbiome response to FOV infection under field conditions, 112 bulk and rhizosphere soil samples were collected from 11 cotton farms (cultivar: Sicot BRF71) located in Macquaire Valley, NSW and St George, QLD (see detailed location in Table S3), which are two of the major cotton-growing regions in Australia where Fusarium wilt is prevalent. Field sampling was done at the flowering stage. At each farm, five healthy and diseased plants from each field were uprooted and gently shaken to remove the soil loosely attached to the root. Similar to glasshouse experiment, disease incidence in field plants were assessed by the presence of brown rings in the cross section of stems. To collect rhizosphere soil samples, soil tightly attached to the root surface was gently brushed off using sterile pencil brushes into sterile plastic bags. To collect the bulk soil samples, five cores of the topsoil (0-20 cm depth) were collected within a radius of 50 cm (1.5 m apart between cotton plants in the field) from each plant before mixing into a sterile plastic bag. Sampling within each field was carried out randomly, but the distance between two samples collected was at least 100 metres. A total number of five samples for each soil compartment (bulk and rhizosphere) were collected in healthy fields, where no disease incidence was reported, and were labelled as *Fusarium*-free samples (FF). In known diseased farms, five healthy (H) and five diseased (D) plants were chosen, and samples were collected for bulk and rhizosphere soil around each plant, as described above. Following the recommended procedure from a previous study (Qiu *et al*., 2020), all samples were stored in sterile bags and buried in ice at the time of sampling and brought to the laboratory. Samples for physiochemical analysis were stored at 4 °C, and samples for molecular analysis were stored at −80 °C until further processing. Soil physiochemical parameters were also measured and described in Table S2.

#### DNA extraction and Illumina MiSeq sequencing

DNA was extracted from frozen soil (~250 mg dry weight) using the DNeasy PowerSoil Pro Kit (Qiagen, Hilden, Germany), following the manufacturer’s instructions. Extracted DNA was quality checked by NanoDrop 2000 (Thermo Fisher Scientific, Waltham, Massachusetts, USA), quantity checked by Qubit Fluorometer (Thermo Fisher Scientific, USA) and PCR checked to confirm the amplifiability following a previous study (Qiu *et al*., 2020).

Amplicon sequencing was carried out targeting the V5-V7 region of 16S rRNA gene for bacterial communities (799F-1193R, Chelius & Triplett, 2001), and ITS2 region (FITS7-ITS4R, Ihrmark *et al*., 2012) for fungal communities. Sequencing was performed at Western Sydney University Next Generation Sequencing (NGS) facility (Richmond, NSW, Australia) using Illumina MiSeq 2 × 300 bp paired end chemistry. All raw sequence data related to this study are available in the NCBI Sequence Read Archive (Assession No. PRJNA770816).

### Microbial community analysis

#### Sequences processing

Raw data obtained from the NGS facility were processed using Mothur (v1.41.3) standard operating procedures (Schloss *et al*., 2009). Briefly, the forward and reverse sequences were merged into contigs. Sequences that contained unidentified bases or had greater than 8 homopolymers were filtered out. For bacterial sequences, an additional step aligning sequences against Silva 16S rRNA gene database version 132 (Pruesse *et al*., 2007) was applied, and unaligned sequences were removed. Refined sequences were pre-clustered (diffs = 1), chimera checked using UCHIME (Edgar *et al*., 2011) and singleton was removed to reduce error (Reeder & Knight, 2009). Bacterial and fungal sequences were then taxonomically classified according to the Silva database version 132 and the UNITE database version 8, respectively, with 60% cut-off confidence. Sequences that match Archaea, cotton mitochondria and chloroplast in bacterial sequences, Alveolata, Metazoa and host ITS regions in fungal sequences were removed. Remaining sequences were clustered into Operational Taxonomic Units (OTUs) at 97% identity and taxonomy was assigned.

#### Statistical analysis and data visualisation

Plant parameters including seed germination, plant height, productivity and disease incidence were analysed in R using ANOVA and visualised using R package “ggplot2” (Lozupone *et al*., 2012). Rarefication of OTU matrices into minimum sample depth and rarefaction curves were conducted using R package “phyloseq”.PERMANOVA was performed using permutational multivariate analysis of variance (Anderson, Marti J., 2001) in PRIMER v. 6 (PRIMER-E, UK) to compare bacterial and fungal communities under different treatments. Similarity matrices were calculated based on Bray-Curtis distances on square-root transformed abundance data to compare the composition and abundances of community structure. Main analyses used 9,999 permutations of residuals under a reduced model (Anderson, M. J., 2001). Pair-wise analyses were performed to compare the differences between treatments with significant interactions based on the main analyses. Alpha and beta diversity were analysed using R package “phyloseq”, and data visualisation plots were generated using Principal Coordinates Analysis (PCoA) using R package “ggplot2”. Linear Discriminant Analysis (LDA) Effective Size (LEfSe) was performed on the Galaxy platform (http://huttenhower.sph.harvard.edu/galaxy/) using one-against-all strategy for multi-class analysis (Segata *et al*., 2011). Phylogenetic trees were generated with Mega X using neighbour-joining method with 1,000 bootstrap before visualised and annotated on iTOL website (Letunic & Bork, 2021).

#### Identification of core members of microbiomes

To identify the persistent microbial species in cotton soil samples and how these species were potentially impacted by FOV, we used a core microbiome approach (Astudillo-García *et al*., 2017) to explore the key members in the cotton rhizosphere microbiome. Briefly, each treatment group (F, FB, and C) from bulk or rhizosphere soil samples, across two time points, were extracted from rarefied bacterial and fungal OTU tables. Following the selection criteria by Hamonts et al. (2018), OTUs that were present in >75% of samples of a particular treatment group were extracted and defined as persistent members of microbiome. Processing of data from field samples was performed in the same way as described above.

#### Co-occurrence network analysis and pathobiome analysis

To infer the bacterial and fungal associations among the OTUs and compare the difference of associations among treatment groups, co-occurrence networks of persistent members of microbiomes were performed using FastSpar (Watts *et al*., 2019), an implementation of network analysis using SparCC algorithm (Friedman & Alm, 2012). Fastspar was run using 50 iterations and 1,000 bootstraps to calculate P-values, with statistically significant correlations (P < 0.05) kept for further analysis. Based on the refined datasets, Gephi (0.9.2) was used to visualise networks using the Fruchterman Reingold layout algorithm (Bastian *et al*., 2009).

To determine the cotton rhizosphere pathobiome, we followed previously reported approaches (Jakuschkin *et al*., 2016; Pauvert *et al*., 2020) using network analysis to subset significant correlations between pathogen and associated microorganisms at the interkingdom scale. To investigate the OTUs closely linked with the pathogen as cotton pathobiome, bacterial and fungal species that correlated with *F. oxysporum* in F and FB treatments were extracted. To confirm the potential biocontrol properties of the pathobiome members that were negatively correlated with *F. oxysporum*, a number of rhizosphere microbes were isolated and characterised from field samples using standard approaches and their activities were tested against the pathogen using plate assays and planta tests (Liu et al., 2021). OTU sequences of candidate pathobiome were matched with sequences from isolated microbes with antagonistic activities against FOV using blastn (McGinnis & Madden, 2004).

## Acknowledgements

Plant microbiome and microbial colonisation work in BKS lab is supported by the Australian Research Council (DP190103714; DP210102081). Field survey work was carried out as a part of project funded by Cotton Research and Development Corporation. We thank the Cotton Info team for their generous help with field survey. JPV thanks to Department of Biotechnology (DBT), India ref. No BT/IACBGF/04/10/2016 for awarding Indo-Australian Career Boosting Gold Fellowships (IACBGF Fellowships 2016-17) for research work in WSU.

## Author’s contributions

ZQ, JPV and BKS designed the experiment; ZQ, JPV and AP performed the experiment; ZQ, BKS, HL, JW, and SK collected the samples; ZQ, HL, JW, BDB and CAD analysed the data; PT, DTT, TW and WC provided the materials; ZQ wrote the manuscript with significant contribution from BKS, and all co-authors reviewed the manuscript.

## Availability of data and materials

All data are publicly available. All raw sequence data related to this study are available in the NCBI Sequence Read Archive (Assession No. PRJNA770816).

## Figure captions

**Figure S1**. Plant data of glasshouse experiment including germination, plant height, productivity and disease incidence throughout the experiment period. C = control treatment (light blue), F = FOV treatment (light red), FB = FOV + biocontrol treatment (light green). There was no significant difference found among different treatments in germination, plant height or productivity.

**Figure S2**. qPCR quantifying FOV load in bulk and the rhizosphere samples. soil.C = bulk soil control (teal), soil.F = bulk soil FOV (yellow), rhizo.C = rhizosphere control treatment (light blue), rhizo.F = rhizosphere FOV treatment (light red), rhizo.FB = rhizosphere FOV + biocontrol treatment (light green).

**Figure S3**. Rarefaction curve for the sequences of soil microbiomes obtained from bacterial 16S rRNA gene sequencing (A) and fungal ITS region sequencing (B) in glasshouse, as well as bacterial 16S rRNA gene sequencing (C) and fungal ITS region sequencing (D) in field samples.

**Figure S4**. Principal Coordinates Analysis (PCoA) plot using Bray-Curtis distance matrix on (A) glasshouse and (B) field samples. Microbial communities in bulk soils (yellow) are different from rhizosphere samples (blue). In glasshouse samples, distinct differences were also found between clay soil (circles) and clay-sandy soil (triangles). In field samples, no significant differences were found in bulk soil (C) between healthy (cyan) and diseased (red) fields, but differences were found in rhizosphere soil (D) between fusarium-free plant (blue) and other plants (diseased: red, healthy: green) in diseased fields. Shapes in (C) and (D) indicate different locations (circles: Macquarie, triangles: St George).

**Figure S5**. Alpha diversity (Chao1, Shannon and Simpson) indices of bacterial (A & C) and fungal (B & D) communities from glasshouse (A & B) and field (C & D) samples. C = control treatment (light blue), F = FOV treatment (light red), FB = FOV + biocontrol treatment (light green), D = diseased plants in diseased field (red), H = healthy plants in diseased field (green), FF = healthy plants in FOV-free field (navy).

## References

Ainsworth TD, Krause L, Bridge T, Torda G, Raina J-B, Zakrzewski M, Gates RD, Padilla-Gamiño JL, Spalding HL, Smith C. 2015. The coral core microbiome identifies rare bacterial taxa as ubiquitous endosymbionts. The ISME journal 9(10): 2261–2274.

Anderson MJ. 2001. A new method for non-parametric multivariate analysis of variance. Austral Ecology 26(1): 32–46.

Anderson MJ. 2001. Permutation tests for univariate or multivariate analysis of variance and regression. Canadian Journal of Fisheries and Aquatic Sciences 58(3): 626–639.

Araujo R, Dunlap C, Barnett S, Franco CMM. 2019. Decoding wheat endosphere-rhizosphere microbiomes in Rhizoctonia solani infested soils challenged by Streptomyces biocontrol agents. Frontiers in plant science 10: 1038.

Arya N, Rana A, Rajwar A, Sahgal M, Sharma A. 2018. Biocontrol efficacy of siderophore producing indigenous Pseudomonas strains against Fusarium Wilt in Tomato. National Academy Science Letters 41(3): 133–136.

Astudillo-García C, Bell JJ, Webster NS, Glasl B, Jompa J, Montoya JM, Taylor MW. 2017. Evaluating the core microbiota in complex communities: a systematic investigation. Environmental Microbiology 19(4): 1450–1462.

Babu S, Seetharaman K, Nandakumar R, Johnson I. 2000. Biocontrol efficacy of Pseudomonas fluorescens against Alternaria solani and tomato leaf blight disease. Annals of Plant Protection Sciences 8(2): 252–254.

Bakker PA, Pieterse CM, de Jonge R, Berendsen RL. 2018. The soil-borne legacy. Cell 172(6): 1178–1180.

Bass D, Stentiford GD, Wang H-C, Koskella B, Tyler CR. 2019. The pathobiome in animal and plant diseases. Trends in Ecology & Evolution 34(11): 996–1008.

Bastian M, Heymann S, Jacomy M 2009. Gephi: an open source software for exploring and manipulating networks. Third international AAAI conference on weblogs and social media.

Benson AK, Kelly SA, Legge R, Ma F, Low SJ, Kim J, Zhang M, Oh PL, Nehrenberg D, Hua K, et al. 2010. Individuality in gut microbiota composition is a complex polygenic trait shaped by multiple environmental and host genetic factors. Proceedings of the National Academy of Sciences 107(44): 18933–18938.

Berendsen RL, Vismans G, Yu K, Song Y, de Jonge R, Burgman WP, Burmølle M, Herschend J, Bakker PA, Pieterse CM. 2018. Disease-induced assemblage of a plant-beneficial bacterial consortium. The ISME journal 12(6): 1496–1507.

Bugbee W. 1970. Vascular response of cotton to infection by Fusarium oxysporum f. sp. vasinfectum. Phytopathology 60(1): 121–123.

Cabrera R, García-López H, Aguirre-von-Wobeser E, Orozco-Avitia JA, Gutiérrez-Saldaña AH. 2020. Amycolatopsis BX17: an actinobacterial strain isolated from soil of a traditional milpa agroecosystem with potential biocontrol against Fusarium graminearum. Biological Control: 104285.

Carrión VJ, Perez-Jaramillo J, Cordovez V, Tracanna V, De Hollander M, Ruiz-Buck D, Mendes LW, Van Ijcken WF, Gomez-Exposito R, Elsayed SS. 2019. Pathogen-induced activation of disease-suppressive functions in the endophytic root microbiome. Science 366(6465): 606–612.

Chaloner TM, Gurr SJ, Bebber DP. 2021. Plant pathogen infection risk tracks global crop yields under climate change. Nature Climate Change 11(8): 710–715.

Chelius M, Triplett E. 2001. The Diversity of Archaea and Bacteria in Association with the Roots of Zea mays L. Microbial Ecology: 252–263.

Chen H, Wu H, Yan B, Zhao H, Liu F, Zhang H, Sheng Q, Miao F, Liang Z. 2018. Core microbiome of medicinal plant Salvia miltiorrhiza seed: a rich reservoir of beneficial microbes for secondary metabolism? International journal of molecular sciences 19(3): 672.

Chen S, Zhang M, Wang J, Lv D, Ma Y, Zhou B, Wang B. 2017. Biocontrol effects of Brevibacillus laterosporus AMCC100017 on potato common scab and its impact on rhizosphere bacterial communities. Biological Control 106: 89–98.

Davis R, Colyer P, Rothrock C, Kochman J. 2006. Fusarium wilt of cotton: population diversity and implications for management. Plant Disease 90(6): 692–703.

Davis R, Moore N, Kochman J. 1996. Characterisation of a population of Fusarium oxysporum f. sp. vasinfectum causing wilt of cotton in Australia. Australian Journal of Agricultural Research 47(7): 1143–1156.

de Jesus Sousa JA, Olivares FL. 2016. Plant growth promotion by streptomycetes: ecophysiology, mechanisms and applications. Chemical and Biological Technologies in Agriculture 3(1): 1–12.

Delgado-Baquerizo M, Guerra CA, Cano-Díaz C, Egidi E, Wang J-T, Eisenhauer N, Singh BK, Maestre FT. 2020. The proportion of soil-borne pathogens increases with warming at the global scale. Nature Climate Change: 1–5.

Develey-Rivière MP, Galiana E. 2007. Resistance to pathogens and host developmental stage: a multifaceted relationship within the plant kingdom. New Phytologist 175(3): 405–416.

Doonan J, Denman S, Pachebat JA, McDonald JE. 2019. Genomic analysis of bacteria in the Acute Oak Decline pathobiome. Microbial genomics 5(1).

Edgar RC, Haas BJ, Clemente JC, Quince C, Knight R. 2011. UCHIME improves sensitivity and speed of chimera detection. Bioinformatics 27(16): 2194–2200.

Elsayed TR, Jacquiod S, Nour EH, Sørensen SJ, Smalla K. 2020. Biocontrol of bacterial wilt disease through complex interaction between tomato plant, antagonists, the indigenous rhizosphere microbiota, and Ralstonia solanacearum. Frontiers in microbiology 10: 2835.

Erlacher A, Cardinale M, Grosch R, Grube M, Berg G. 2014. The impact of the pathogen Rhizoctonia solani and its beneficial counterpart Bacillus amyloliquefaciens on the indigenous lettuce microbiome. Frontiers in Microbiology 5: 175.

Essarioui A, LeBlanc N, Kistler HC, Kinkel LL. 2017. Plant community richness mediates inhibitory interactions and resource competition between Streptomyces and Fusarium populations in the rhizosphere. Microbial Ecology 74(1): 157–167.

Fierer N, Jackson JA, Vilgalys R, Jackson RB. 2005. Assessment of soil microbial community structure by use of taxon-specific quantitative PCR assays. Applied and Environmental Microbiology 71(7): 4117–4120.

Friedman J, Alm EJ. 2012. Inferring correlation networks from genomic survey data. PLoS Comput Biol 8(9): e1002687.

Gajbhiye A, Rai AR, Meshram SU, Dongre A. 2010. Isolation, evaluation and characterization of Bacillus subtilis from cotton rhizospheric soil with biocontrol activity against Fusarium oxysporum. World Journal of Microbiology and Biotechnology 26(7): 1187–1194.

Göre ME, Caner ÖK, Altin N, Aydin MH, Erdogan O, Filizer F, Büyükdögerlioglu A. 2009. Evaluation of cotton cultivars for resistance to pathotypes of Verticillium dahliae. Crop Protection 28(3): 215–219.

Goudjal Y, Zamoum M, Sabaou N, Mathieu F, Zitouni A. 2016. Potential of endophytic Streptomyces spp. for biocontrol of Fusarium root rot disease and growth promotion of tomato seedlings. Biocontrol Science and Technology 26(12): 1691–1705.

Gu Y, Wei Z, Wang X, Friman V-P, Huang J, Wang X, Mei X, Xu Y, Shen Q, Jousset A. 2016. Pathogen invasion indirectly changes the composition of soil microbiome via shifts in root exudation profile. Biology and Fertility of Soils 52(7): 997–1005.

Hamonts K, Trivedi P, Garg A, Janitz C, Grinyer J, Holford P, Botha FC, Anderson IC, Singh BK. 2018. Field study reveals core plant microbiota and relative importance of their drivers. Environmental Microbiology 20(1): 124–140.

Hoffman MT, Arnold AE. 2010. Diverse bacteria inhabit living hyphae of phylogenetically diverse fungal endophytes. Applied and Environmental Microbiology 76(12): 4063–4075.

Hollomon DW. 2015. Fungicide resistance: facing the challenge-a review. Plant protection science 51(4): 170–176.

Hu J, Wei Z, Friman V-P, Gu S-h, Wang X-f, Eisenhauer N, Yang T-j, Ma J, Shen Q-r, Xu Y-c. 2016. Probiotic diversity enhances rhizosphere microbiome function and plant disease suppression. MBio 7(6): e01790–01716.

Ihrmark K, Bödeker I, Cruz-Martinez K, Friberg H, Kubartova A, Schenck J, Strid Y, Stenlid J, Brandström-Durling M, Clemmensen KE. 2012. New primers to amplify the fungal ITS2 region–evaluation by 454-sequencing of artificial and natural communities. FEMS Microbiology Ecology 82(3): 666–677.

Jakuschkin B, Fievet V, Schwaller L, Fort T, Robin C, Vacher C. 2016. Deciphering the pathobiome: intra-and interkingdom interactions involving the pathogen Erysiphe alphitoides. Microbial Ecology 72(4): 870–880.

Jangir M, Pathak R, Sharma S, Sharma S. 2018. Biocontrol mechanisms of Bacillus sp., isolated from tomato rhizosphere, against Fusarium oxysporum f. sp. lycopersici. Biological Control 123: 60–70.

Johnson ET, Bowman MJ, Dunlap CA. 2020. Brevibacillus fortis NRS-1210 produces edeines that inhibit the in vitro growth of conidia and chlamydospores of the onion pathogen Fusarium oxysporum f. sp. cepae. Antonie Van Leeuwenhoek 113(7): 973–987.

Kaushal M, Swennen R, Mahuku G. 2020. Unlocking the Microbiome Communities of Banana (Musa spp.) under Disease Stressed (Fusarium wilt) and Non-Stressed Conditions. Microorganisms 8(3): 443.

Khan N, Maymon M, Hirsch AM. 2017. Combating Fusarium infection using Bacillus-based antimicrobials. Microorganisms 5(4): 75.

Krezalek MA, DeFazio J, Zaborina O, Zaborin A, Alverdy JC. 2016. The shift of an intestinal “microbiome” to a “pathobiome” governs the course and outcome of sepsis following surgical injury. Shock (Augusta, Ga.) 45(5): 475.

Kulmatiski A, Beard KH. 2011. Long-term plant growth legacies overwhelm short-term plant growth effects on soil microbial community structure. Soil Biology and Biochemistry 43(4): 823–830.

Kwak M-J, Kong HG, Choi K, Kwon S-K, Song JY, Lee J, Lee PA, Choi SY, Seo M, Lee HJ. 2018. Rhizosphere microbiome structure alters to enable wilt resistance in tomato. Nature Biotechnology 36(11): 1100–1109.

Lamb EG, Kennedy N, Siciliano SD. 2011. Effects of plant species richness and evenness on soil microbial community diversity and function. Plant and Soil 338(1-2): 483–495.

Lane D. 1991. 16S/23S rRNA sequencing. Nucleic acid techniques in bacterial systematics: 115–175.

Lay C-Y, Bell TH, Hamel C, Harker KN, Mohr R, Greer CW, Yergeau É, St-Arnaud M. 2018. Canola root–associated microbiomes in the Canadian Prairies. Frontiers in microbiology 9: 1188.

Lemanceau P, Blouin M, Muller D, Moënne-Loccoz Y. 2017. Let the core microbiota be functional. Trends in Plant Science 22(7): 583–595.

Leoni C, Piancone E, Sasanelli N, Bruno GL, Manzari C, Pesole G, Ceci LR, Volpicella M. 2020. Plant Health and Rhizosphere Microbiome: Effects of the Bionematicide Aphanocladium album in Tomato Plants Infested by Meloidogyne javanica. Microorganisms 8(12): 1922.

Letunic I, Bork P. 2021. Interactive Tree Of Life (iTOL) v5: an online tool for phylogenetic tree display and annotation. Nucleic Acids Research 49(W1): W293–W296.

Liu H, Brettell LE, Qiu Z, Singh BK. 2020. Microbiome-mediated stress resistance in plants. Trends in Plant Science.

Liu H, Li J, Carvalhais LC, Percy CD, Prakash Verma J, Schenk PM, Singh BK. 2021a. Evidence for the plant recruitment of beneficial microbes to suppress soil - borne pathogens. New Phytologist 229(5): 2873–2885.

Liu H, Qiu Z, Ye J, Verma JP, Li J, Singh BK. 2021b. Effective colonisation by a bacterial synthetic community promotes plant growth and alters soil microbial community. Journal of Sustainable Agriculture and Environment.

Lozupone CA, Stombaugh JI, Gordon JI, Jansson JK, Knight R. 2012. Diversity, stability and resilience of the human gut microbiota. Nature 489(7415): 220–230.

Lucas JA, Hawkins NJ, Fraaije BA. 2015. The evolution of fungicide resistance. Advances in Applied Microbiology 90: 29–92.

McGinnis S, Madden TL. 2004. BLAST: at the core of a powerful and diverse set of sequence analysis tools. Nucleic Acids Research 32(suppl_2): W20–W25.

Misk A, Franco C. 2011. Biocontrol of chickpea root rot using endophytic actinobacteria. BioControl 56(5): 811–822.

Padda KP, Puri A, Chanway CP 2017. Paenibacillus polymyxa: A prominent biofertilizer and biocontrol agent for sustainable agriculture. Agriculturally important microbes for sustainable agriculture: Springer, 165–191.

Palaniyandi SA, Yang SH, Zhang L, Suh J-W. 2013. Effects of actinobacteria on plant disease suppression and growth promotion. Applied Microbiology and Biotechnology 97(22): 9621–9636.

Pauvert C, Fort T, Calonnec A, d’Arcier JF, Chancerel E, Massot M, Chiquet J, Robin S, Bohan DA, Vallance J. 2020. Microbial association networks give relevant insights into plant pathobiomes. bioRxiv.

Pruesse E, Quast C, Knittel K, Fuchs BM, Ludwig W, Peplies J, Glöckner FO. 2007. SILVA: a comprehensive online resource for quality checked and aligned ribosomal RNA sequence data compatible with ARB. Nucleic Acids Research 35.

Qiu Z, Egidi E, Liu H, Kaur S, Singh BK. 2019. New frontiers in agriculture productivity: Optimised microbial inoculants and in situ microbiome engineering. Biotechnology Advances.

Qiu Z, Wang J, Delgado-Baquerizo M, Trivedi P, Egidi E, Chen Y-M, Zhang H, Singh BK. 2020. Plant microbiomes: do different preservation approaches and primer sets alter our capacity to assess microbial diversity and community composition? Frontiers in plant science 11: 993.

Ramakers C, Ruijter JM, Deprez RHL, Moorman AF. 2003. Assumption-free analysis of quantitative real-time polymerase chain reaction (PCR) data. Neuroscience Letters 339(1): 62–66.

Ramakrishna N, Lacey J, Smith J. 1991. Effect of surface sterilization, fumigation and gamma irradiation on the microflora and germination of barley seeds. International Journal of Food Microbiology 13(1): 47–54.

Reeder J, Knight R. 2009. The ‘rare biosphere’: a reality check. Nature Methods 6(9): 636–637.

Rybakova D, Mancinelli R, Wikström M, Birch-Jensen A-S, Postma J, Ehlers R-U, Goertz S, Berg G. 2017. The structure of the Brassica napus seed microbiome is cultivar-dependent and affects the interactions of symbionts and pathogens. Microbiome 5(1): 104.

Saravanakumar K, Li Y, Yu C, Wang Q-q, Wang M, Sun J, Gao J-x, Chen J. 2017. Effect of Trichoderma harzianum on maize rhizosphere microbiome and biocontrol of Fusarium Stalk rot. Scientific reports 7(1): 1–13.

Schlatter D, Kinkel L, Thomashow L, Weller D, Paulitz T. 2017. Disease suppressive soils: new insights from the soil microbiome. Phytopathology 107(11): 1284–1297.

Schlatter DC, Yin C, Hulbert S, Paulitz TC. 2020. Core rhizosphere microbiomes of dryland wheat are influenced by location and land use history. Applied and Environmental Microbiology 86(5).

Schloss PD, Westcott SL, Ryabin T, Hall JR, Hartmann M, Hollister EB, Lesniewski RA, Oakley BB, Parks DH, Robinson CJ, et al. 2009. Introducing mothur: Open-Source, Platform-Independent, Community-Supported Software for Describing and Comparing Microbial Communities. Applied and Environmental Microbiology 75(23): 7537–7541.

Segata N, Izard J, Waldron L, Gevers D, Miropolsky L, Garrett WS, Huttenhower C. 2011. Metagenomic biomarker discovery and explanation. Genome biology 12(6): 1–18.

Shade A, Handelsman J. 2012. Beyond the Venn diagram: the hunt for a core microbiome. Environmental Microbiology 14(1): 4–12.

Shen Z, Penton CR, Lv N, Xue C, Yuan X, Ruan Y, Li R, Shen Q. 2018. Banana Fusarium Wilt Disease Incidence Is Influenced by Shifts of Soil Microbial Communities Under Different Monoculture Spans. Microbial Ecology 75(3): 739–750.

Singh BK, Trivedi P, Egidi E, Macdonald CA, Delgado-Baquerizo M. 2020. Crop microbiome and sustainable agriculture. Nature Reviews Microbiology 18(11): 601–602.

Sun D, Zhuo T, Hu X, Fan X, Zou H. 2017. Identification of a Pseudomonas putida as biocontrol agent for tomato bacterial wilt disease. Biological Control 114: 45–50.

Sweet M, Burian A, Fifer J, Bulling M, Elliott D, Raymundo L. 2019. Compositional homogeneity in the pathobiome of a new, slow-spreading coral disease. Microbiome 7(1): 1–14.

Sweet MJ, Bulling MT. 2017. On the importance of the microbiome and pathobiome in coral health and disease. Frontiers in Marine Science 4: 9.

Szczech M, Shoda M. 2006. The Effect of mode of application of Bacillus subtilis RB14-C on its efficacy as a biocontrol agent against Rhizoctonia solani. Journal of Phytopathology 154(6): 370–377.

Thomas T, Moitinho-Silva L, Lurgi M, Björk JR, Easson C, Astudillo-García C, Olson JB, Erwin PM, López-Legentil S, Luter H. 2016. Diversity, structure and convergent evolution of the global sponge microbiome. Nature communications 7(1): 1–12.

Trivedi P, Delgado-Baquerizo M, Jeffries TC, Trivedi C, Anderson IC, Lai K, McNee M, Flower K, Pal Singh B, Minkey D. 2017a. Soil aggregation and associated microbial communities modify the impact of agricultural management on carbon content. Environmental Microbiology 19(8): 3070–3086.

Trivedi P, Leach JE, Tringe SG, Sa T, Singh BK. 2020. Plant–microbiome interactions: from community assembly to plant health. Nature Reviews Microbiology 18(11): 607–621.

Trivedi P, Schenk PM, Wallenstein MD, Singh BK. 2017b. Tiny microbes, big yields: enhancing food crop production with biological solutions. Microbial biotechnology 10(5): 999–1003.

Tufts DM, Sameroff S, Tagliafierro T, Jain K, Oleynik A, VanAcker MC, Diuk-Wasser MA, Lipkin WI, Tokarz R. 2020. A metagenomic examination of the pathobiome of the invasive tick species, Haemaphysalis longicornis, collected from a New York City borough, USA. Ticks and Tick-borne Diseases 11(6): 101516.

Turnbaugh PJ, Ley RE, Hamady M, Fraser-Liggett CM, Knight R, Gordon JI. 2007. The Human Microbiome Project. Nature 449(7164): 804–810.

Ulloa M, Hutmacher RB, Schramm T, Ellis ML, Nichols R, Roberts PA, Wright SD. 2020. Sources, selection and breeding of Fusarium wilt (Fusarium oxysporum f. sp. vasinfectum) race 4 (FOV4) resistance in Upland (Gossypium hirsutum L.) cotton. Euphytica 216(7): 1–18.

Vandenkoornhuyse P, Quaiser A, Duhamel M, Le Van A, Dufresne A. 2015. The importance of the microbiome of the plant holobiont. New Phytologist 206(4): 1196–1206.

Vayssier-Taussat M, Albina E, Citti C, Cosson JF, Jacques M-A, Lebrun M-H, Le Loir Y, Ogliastro M, Petit M-A, Roumagnac P. 2014. Shifting the paradigm from pathogens to pathobiome: new concepts in the light of meta-omics. Frontiers in cellular and infection microbiology 4: 29.

Wachowska U, Irzykowski W, Jędryczka M, Stasiulewicz-Paluch AD, Głowacka K. 2013. Biological control of winter wheat pathogens with the use of antagonistic Sphingomonas bacteria under greenhouse conditions. Biocontrol Science and Technology 23(10): 1110–1122.

Wang T, Hao Y, Zhu M, Yu S, Ran W, Xue C, Ling N, Shen Q. 2019. Characterizing differences in microbial community composition and function between Fusarium wilt diseased and healthy soils under watermelon cultivation. Plant and Soil 438(1): 421–433.

Watts SC, Ritchie SC, Inouye M, Holt KE. 2019. FastSpar: rapid and scalable correlation estimation for compositional data. Bioinformatics 35(6): 1064–1066.

Wei Z, Gu Y, Friman V-P, Kowalchuk GA, Xu Y, Shen Q, Jousset A. 2019. Initial soil microbiome composition and functioning predetermine future plant health. Science advances 5(9): eaaw0759.

Wei Z, Hu J, Yin S, Xu Y, Jousset A, Shen Q, Friman V-P. 2018. Ralstonia solanacearum pathogen disrupts bacterial rhizosphere microbiome during an invasion. Soil Biology and Biochemistry 118: 8–17.

Wei Z, Yang T, Friman V-P, Xu Y, Shen Q, Jousset A. 2015. Trophic network architecture of root-associated bacterial communities determines pathogen invasion and plant health. Nature communications 6(1): 1–9.

Xiong C, He JZ, Singh BK, Zhu YG, Wang JT, Li PP, Zhang QB, Han LL, Shen JP, Ge AH. 2021a. Rare taxa maintain the stability of crop mycobiomes and ecosystem functions. Environmental Microbiology 23(4): 1907–1924.

Xiong C, Zhu YG, Wang JT, Singh B, Han LL, Shen JP, Li PP, Wang GB, Wu CF, Ge AH. 2021b. Host selection shapes crop microbiome assembly and network complexity. New Phytologist 229(2): 1091–1104.

Xu J, Zhang Y, Zhang P, Trivedi P, Riera N, Wang Y, Liu X, Fan G, Tang J, Coletta-Filho HD. 2018. The structure and function of the global citrus rhizosphere microbiome. Nature communications 9(1): 1–10.

Yang L, Xie J, Jiang D, Fu Y, Li G, Lin F. 2008. Antifungal substances produced by Penicillium oxalicum strain PY-1—potential antibiotics against plant pathogenic fungi. World Journal of Microbiology and Biotechnology 24(7): 909–915.

Zambounis A, Paplomatas E, Tsaftaris A. 2007. Intergenic spacer–RFLP analysis and direct quantification of Australian Fusarium oxysporum f. sp. vasinfectum isolates from soil and infected cotton tissues. Plant Disease 91(12): 1564–1573.

