## Supplementary material for "Response of the plant core microbiome to *Fusarium oxysporum* infection and identification of the pathobiome": Supplementary.docx

**Supplementary Information**

**Preparation of microbial isolates**

Candidates of bacterial isolates were sourced from BKS lab culture collections, where bacterial strains were isolated from rhizosphere soil collected from cotton field in in Narrabri, NSW, Australia. A total number of 45 bacterial isolates as glycerol stocks were revived on Luria-Bertani (LB) agar plates by streaking methods and incubated at 28 ºC for 24 h. Single colonies were subcultured in liquid LB media till exponential increasing stage before further analyses. Fungal pathogen *Fusarium oxysporum* f.sp. *vasinfactum* strain FOV 294-2 (thereafter FOV) was also sourced from BKS lab culture collections, which was originally collected from FOV-infected cotton plants. FOV was firstly revived on LB agar at 28 ºC for 24 h, before subculturing on new agar plates for antagonistic assay.

**Antagonistic and synergistic assay**

For antagonistic assay, a circular mycelial disc of the actively growing FOV was placed at the centre of PDA plate along with four bacterial isolates streaked evenly on the side of the plate. Antagonistic plates were incubated at 28 ºC until distinctive inhibition patterns can be observed. Antifungal activity of each bacterial isolate was then evaluated by measuring the size of the colony. Of the 45 revived bacterial isolates, 32 isolates showed antagonistic properties against FOV (Table S1).

For synergistic assay, each pair of bacterial strains were inoculated on one agar plate following the method described by Berendsen et al. (2018). Isolates without antagonistic mechanism against other bacterial strains were selected as biocontrol agents. A total number of seven strains with antifungal activity against FOV were selected for further characterisation step.

**Characterisation of microbial strains**

Seven selected bacterial strains (Table S1) and FOV were characterised by Sanger sequencing (Qiagen, Hilden, Germany). For each microbial strain, a loopful of bacteria/fungi from the agar plate was collected and transferred into an Eppendorf tube with 200 µl TE buffer (10 mM Tris-HCl, 1mM EDTA, pH 8.0), vortexed to mix thoroughly. After boiled for 5 min, 1 µl of microbial solution were applied as template in 25 µl PCR system. PCR was performed using primer pairs (27F 5'-AGAGTTTGATCCTGGCTCAG-3', 1492R 5'-GGTTACCTTGTTACGACTT-3') (Lane, 1991) targeting 16S rRNA gene on bacteria, respectively. The PCR was conducted as follows: 94 ºC 5 min for denaturation, followed by 30 cycles of amplification (94 ºC 30 s, 55 ºC 30 s, 72 ºC 90 s), then 72 ºC 7 min for elongation. PCR products were cleaned using QIAquick PCR Purification Kit (Qiagen, Hilden, Germany) before continuing with Sanger sequencing.

Microbial strains were characterised using sequences obtained from the sequencing with BLAST at NCBI website ([https://blast.ncbi.nlm.nih.gov/Blast.cgi](about:blank)).

**References**

**Berendsen RL, Vismans G, Yu K, Song Y, de Jonge R, Burgman WP, Burmølle M, Herschend J, Bakker PA, Pieterse CM. 2018.** Disease-induced assemblage of a plant-beneficial bacterial consortium. *The ISME journal* **12**(6): 1496-1507.

**Lane D. 1991.** 16S/23S rRNA sequencing. *Nucleic acid techniques in bacterial systematics*: 115-175.
