## Supplementary figures and images for "Response of the plant core microbiome to *Fusarium oxysporum* infection and identification of the pathobiome"

### Figure S1 - plant data.pdf

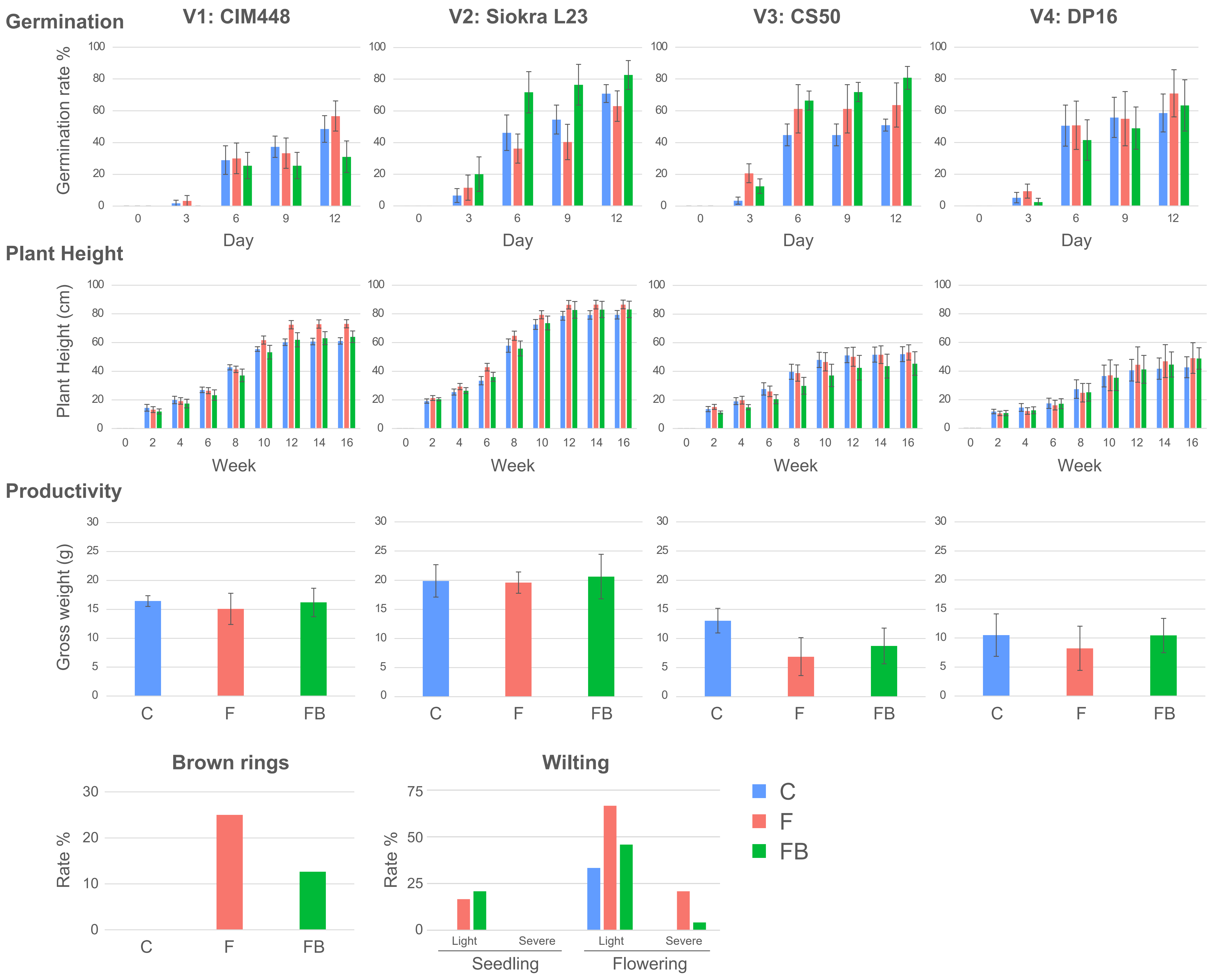

### Figure S2 - qPCR.pdf

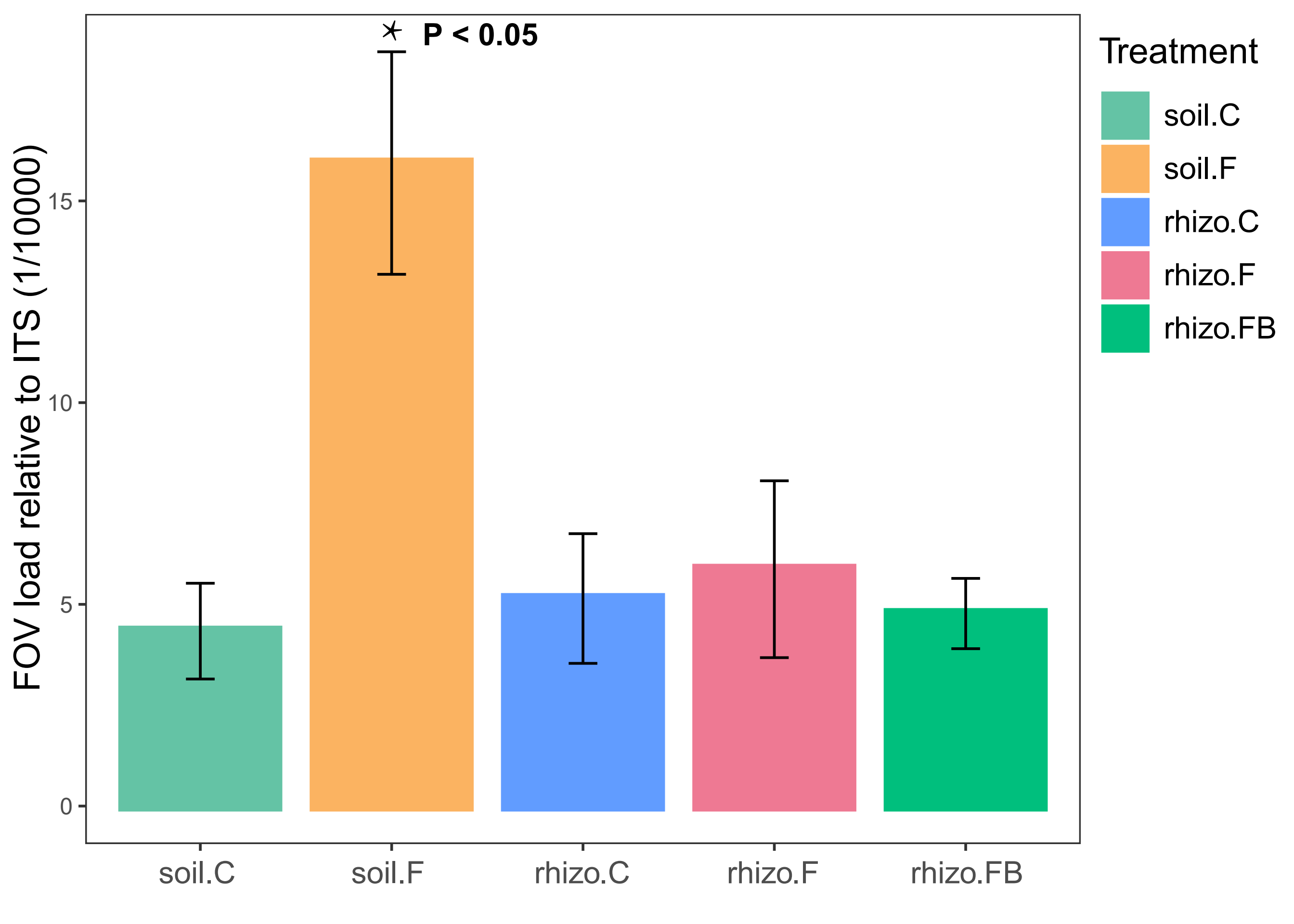

### Figure S3 - Rarefaction curve.pdf

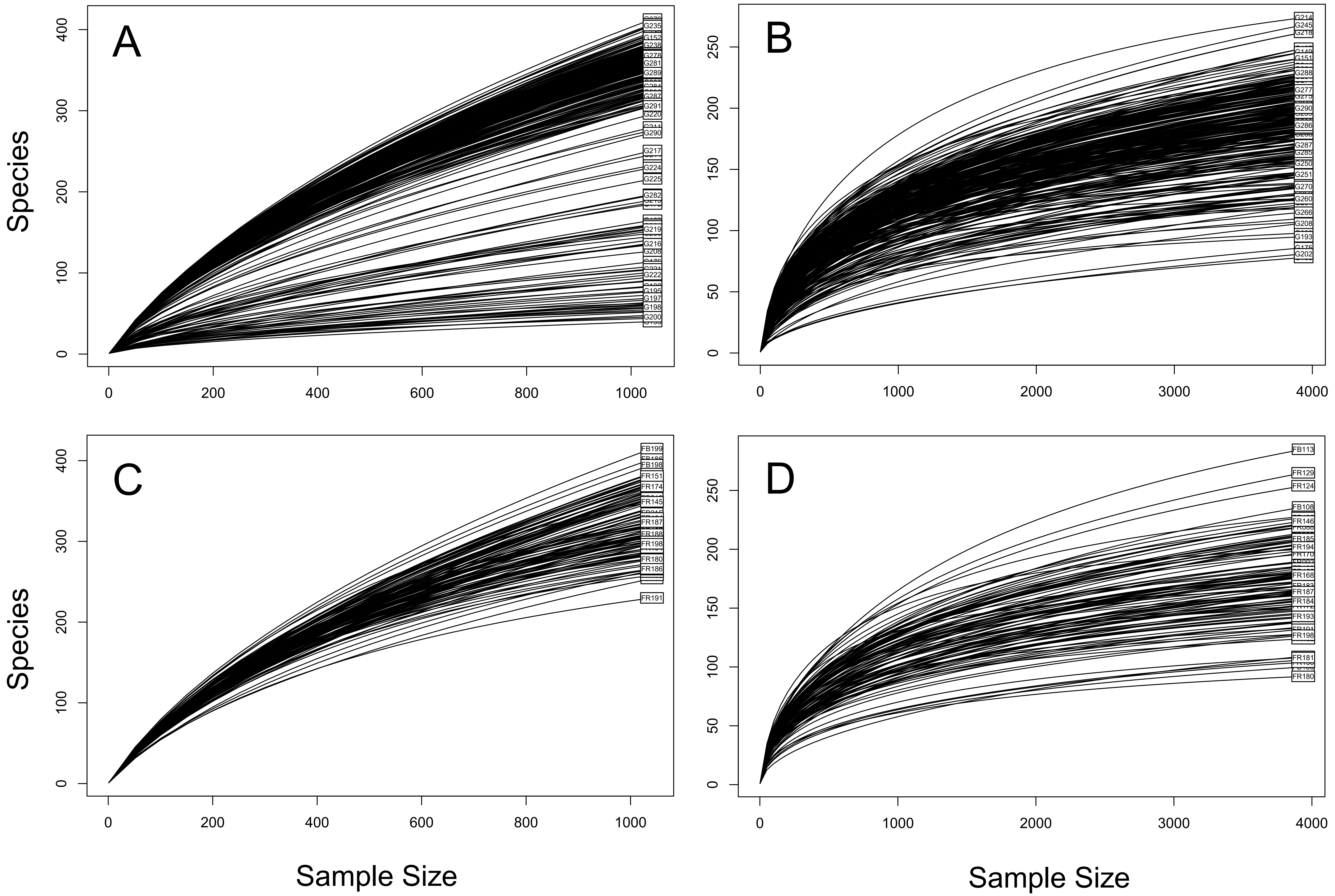

### Figure S4 - PCoA soil.pdf

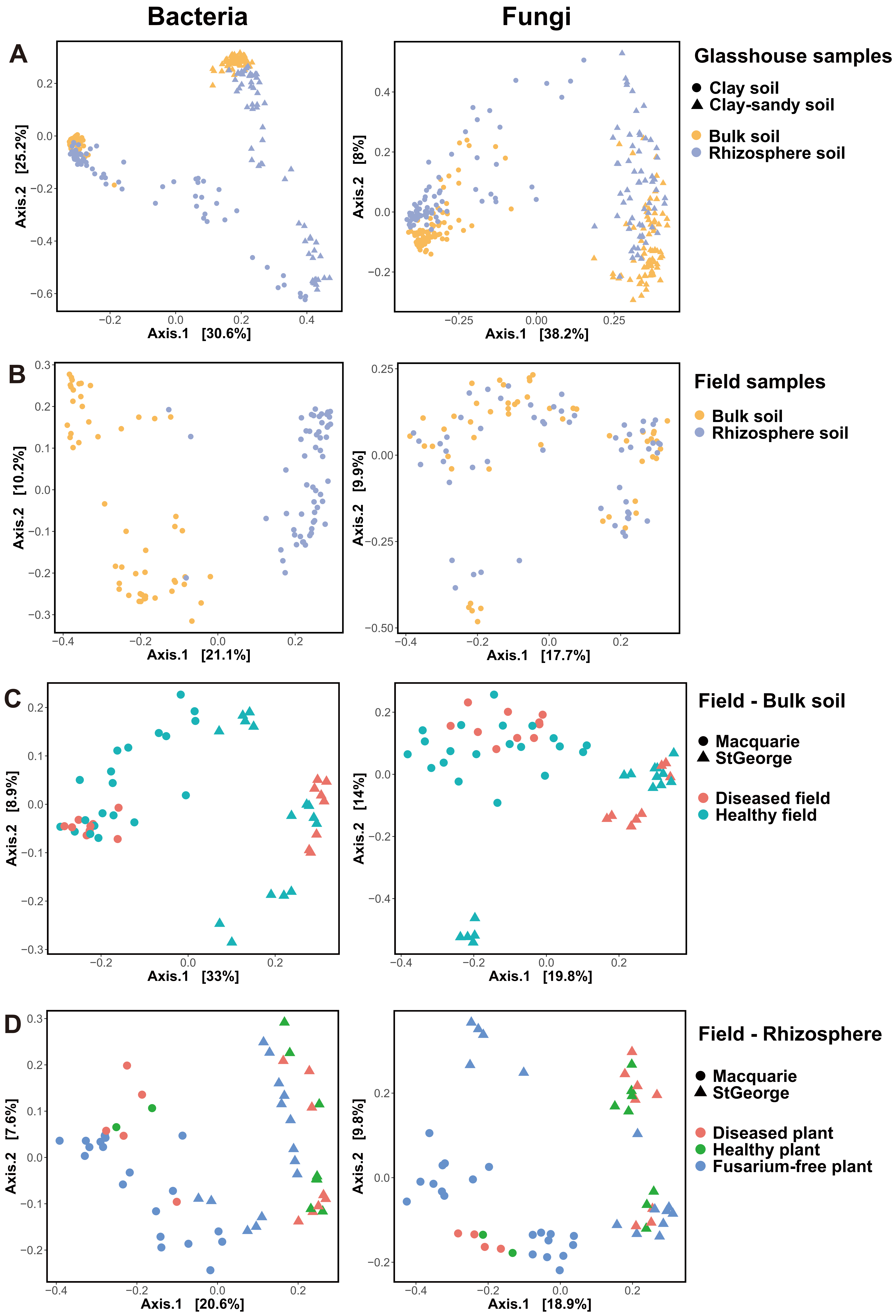

### Figure S5 - overall alpha diversity.pdf

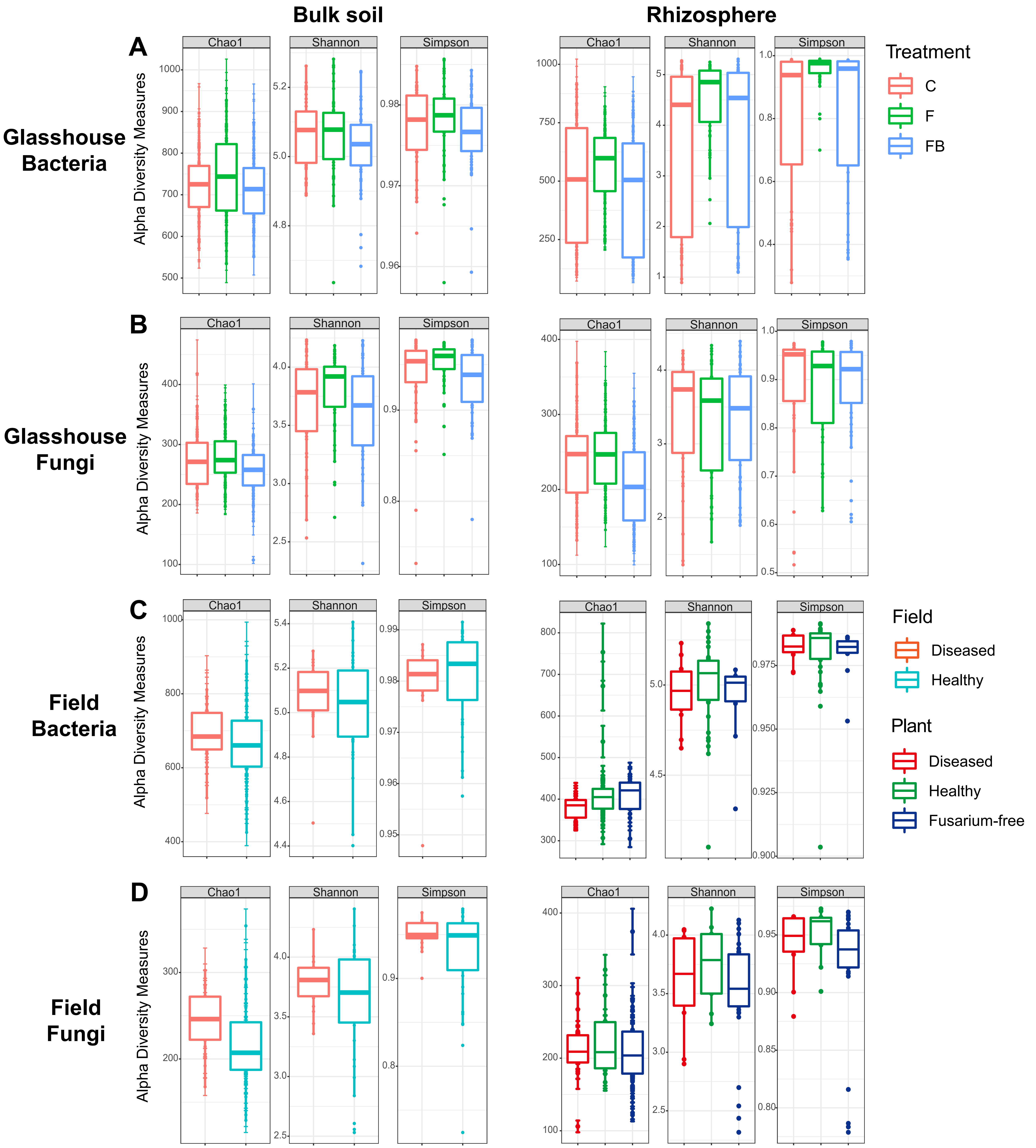
